# A strategy to suppress STAT1 signalling conserved in pathogenic poxviruses and paramyxoviruses

**DOI:** 10.1101/2021.07.17.452491

**Authors:** Callum Talbot-Cooper, Teodors Pantelejevs, John P. Shannon, Christian R. Cherry, Marcus T. Au, Marko Hyvönen, Heather D. Hickman, Geoffrey L. Smith

**Affiliations:** Department of Pathology, University of Cambridge, Tennis Court Road, Cambridge, CB2 1QP; Department of Biochemistry, University of Cambridge, 80 Tennis Court Road, Cambridge, CB2 1GA; Viral Immunity and Pathogenesis Unit, Laboratory of Clinical Immunology and Microbiology, National Institute of Allergy and Infectious Diseases, National Institutes of Health, Bethesda, Maryland 20852, USA

## Abstract

The induction of interferon-stimulated genes by signal transducer and activator of transcription (STAT) proteins, is a critical host defence to fight virus infections. Here, a highly expressed poxvirus protein 018 is shown to inhibit IFN-induced signalling by binding the SH2 domain of STAT1 to prevent STAT1 association with an activated IFN receptor. Despite the presence of additional inhibitors of IFN-induced signalling, a poxvirus lacking 018 was attenuated in mice. The 2.0 Å crystal structure of the 018:STAT1 complex reveals a mechanism for a high-affinity, pTyr-independent mode of binding to an SH2 domain. Furthermore, the STAT1 binding motif of 018 shows sequence similarity to the STAT1-binding proteins from Nipah virus, which like 018, block the association of STAT1 with an IFN receptor. Taken together, these results provide detailed mechanistic insight into a potent mode of STAT1 antagonism, found to exist in genetically diverse virus families.

## Introduction

Interferons (IFNs) activate signal transduction pathways to upregulate IFN-stimulated genes (ISGs) that inhibit virus replication and spread (Schneider et al., 2014). Signal transduction is mediated by signal transducers of transcription (STAT) proteins STAT1 and STAT2, which, in an unstimulated state, exist as latent unphosphorylated hetero (U-STAT1-U-STAT2) or homodimers (U-STAT1) (Mao et al., 2005; Wang et al., 2021). IFNs bind their cognate receptors to activate receptor-associated kinases that in turn phosphorylate receptor tails creating a docking site for STAT SH2 domains. At the receptors, STATs are phosphorylated (pSTAT) and undergo dimer rearrangement from an anti-parallel to an activated parallel conformation, mediated by a reciprocal pTyr:SH2 interaction between two pSTATs (Wenta et al., 2008).

Type I IFNs (IFN-I) signal through the IFNα/β receptor (IFNAR) to activate kinases that phosphorylate STAT1 and 2. The pSTAT1:STAT2 heterodimer associates with IRF9 to form a trimeric complex known as IFN-stimulated gene factor 3 (ISGF3) (Rengachari et al., 2018). Type II IFN (IFN-II), of which IFNγ is the sole member, signals through the IFNγ receptor (IFNGR) and activates kinases that phosphorylate STAT1 only. The pSTAT1 homodimer is known as the gamma-activated factor (GAF). Nuclear ISGF3 and GAF drive the transcription of ISGs with IFN-stimulated responsive element (ISRE) or gamma-activated sequence (GAS) promoters, respectively (Aaronson and Horvath, 2002).

To overcome the anti-viral activities induced by IFNs, viruses have evolved numerous strategies to antagonise host IFN-pathways, for reviews see (García-Sastre, 2017; Randall and Goodbourn, 2008). Given the importance of viral-mediated IFN-signalling antagonism for productive infection, mechanistic insight into these strategies can guide novel anti-viral therapeutic approaches.

Poxviruses are large, cytoplasmic DNA viruses. Vaccinia virus (VACV) is the prototypic poxvirus, the vaccine used to eradicate smallpox and an excellent model to study host-pathogen interactions. VACV encodes about 200 proteins of which it is estimated that >1/3 modulate host immune responses, including proteins that target IFN-induced signalling pathways (Smith et al., 2013, 2018). VACV proteins B18 and B8 act as soluble IFN receptors that bind IFN-I and IFN-II, respectively (Alcamí and Smith, 1995; Colamonici et al., 1995; Mossman et al., 1995; Symons et al., 1995). At the intracellular level, the viral phosphatase vH1 dephosphorylates STAT1 (Koksal et al., 2009; Najarro et al., 2001), whilst protein C6 inhibits IFN-I signalling in the nucleus (Stuart et al., 2016).

Here we show that an uncharacterised viral protein (018), encoded by gene VACWR018 of VACV strain Western Reserve (WR), binds directly to the SH2 domain of STAT1 and competes with a phosphorylated IFN receptor to prevent STAT1 receptor association and subsequent STAT1 phosphorylation. A VACV lacking 018 was attenuated in mice and induced enhanced innate immune signalling, demonstrating the *in vivo* importance of this inhibitor of IFN-induced signalling for poxviruses. The crystal structure of 018 complexed with STAT1 was determined to 2.0 Å. This revealed a key contact that enables 018 to bind STAT1 and STAT4 selectively, and a non-canonical SH2 binding mode, whereby 018 occupies the SH2 domain in a pTyr pocket-independent manner with high affinity. The 018 STAT1-binding motif is identical in variola virus and monkeypox virus orthologues of 018. In addition, it shares remarkable similarity to STAT1-binding regions of V/W and P proteins from Nipah virus (NiV), a highly pathogenic paramyxovirus. Like 018, we show that the minimal STAT1 binding region of NiV-V protein can compete with a phosphorylated IFN receptor to bind STAT1. This study reveals a conserved mechanism for targeting STAT1 utilised by members of the poxvirus and paramyxovirus families, to subvert cellular anti-viral responses.

## Results

The 018 open reading frame (ORF) from VACV WR (gene VACWR018) is transcribed early during infection and is one of the most abundant viral transcripts (Assarsson et al., 2008; Wennier et al., 2013; Yang et al., 2015). Protein 018 is highly conserved within the orthopoxvirus genus including human pathogens cowpox virus, monkeypox virus and variola virus, the causative agent of smallpox (**Figure S1**). The 018 ORF is also highly conserved in ancient variola viruses dating from the Viking age (Mühlemann et al., 2020).

As 018 is expressed early during infection, is highly abundant, and encoded within a region of the genome known to harbour immunomodulators (Gubser et al., 2004), we explored if 018 modulates anti-viral immunity.

### Vaccinia protein 018 inhibits IFN-induced signalling

To test if 018 modulates anti-viral immunity, reporter plasmids were used that express a luciferase (Luc) gene upon activation of specific anti-viral signalling pathways. Activation of IRF3, NF-kB and AP-1 pathways that induce IFNs (specifically IFNβ) were measured using an IFNβ-Luc reporter after stimulation with Sendai virus (SeV), the prototypic paramyxovirus. Downstream, IFN-induced pathways were measured using ISRE or GAS-Luc reporters after stimulation with IFN-I (IFNα) or IFN-II (IFNγ), respectively.

TAP-tagged (2 X Strep, 1 X FLAG epitope) 018 inhibited pathway activation induced by IFN-I and II (**Figure 1A, B**), whereas it had little effect on the activation of IFNβ-Luc (**Figure 1C**).

**Figure 1.**
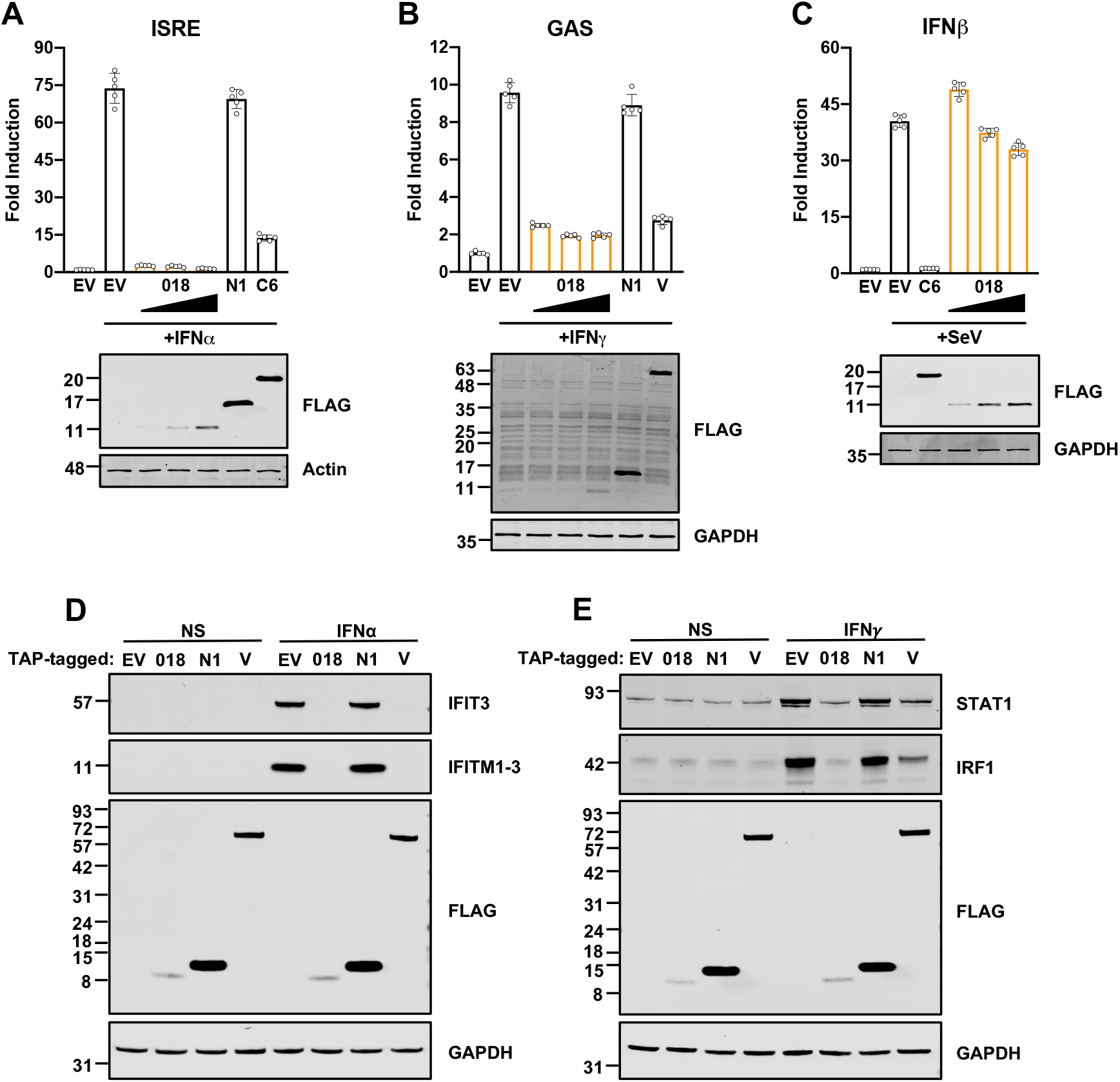
Vaccinia protein 018 inhibits IFN-induced signalling. **(A, C)** HEK 293T cells or **(B)** HeLa cells were transfected with reporter plasmids ISRE-Luc **(A)**, GAS-Luc **(B),** or IFNβ-Luc **(C)**, plus *TK-Renilla* and vectors expressing proteins indicated fused to a TAP-tag or empty vector (EV). Cells were stimulated with IFNα (1000 U/mL) **(A)**, IFNγ (25 ng/mL) **(B)** or SeV **(C)**, for 6 **(A)**, 8 **(B)** or 24 h **(C)** and then luciferase activity was measured. Means ± SD (n=5 per condition) are shown. **(D-E)** T-REx 293 cells were induced with doxycycline (dox, 100 ng/mL) to express indicated proteins. Cells were non-stimulated (NS) or stimulated with IFNα (1000 U/mL) **(D)** or IFNγ (25 ng/mL) **(E)** for 24 h and lysates were analysed by immunoblotting. Immunoblots were stained against proteins/epitope indicated (**A-E**). Data for (**A-C**) and (**D-E**) are representative of three or two individual experiments, respectively.

NiV-V (Rodriguez et al., 2002) and VACV protein C6 (Stuart et al., 2016; Unterholzner et al., 2011) were used as positive controls, whereas VACV protein N1 (Maluquer de Motes et al., 2011) was used as a negative control.

Next, the effect of 018 on expression of endogenous ISGs was tested. T-REx 293 cell lines that inducibly expressed TAP-tagged 018 or controls (EV, TAP-tagged N1 or NiV-V) were stimulated with IFN-I or -II and representative ISGs were analysed by immunoblotting. Stimulation with IFN led to an increase in ISG levels in cells expressing either EV, or N1 (**Figure 1D, E**). In contrast, 018 blocked upregulation of ISGs after stimulation with either IFN-I or IFN-II **(Figure 1D, E**). These data show 018 is a potent inhibitor of IFN-I and -II-induced signalling.

### Phosphorylation of STAT1 at Tyr 701 is blocked by 018

To assess if 018 inhibited STAT translocation into the nucleus, HeLa cells were transfected with TAP-tagged 018 or controls and then stimulated with IFN-I or -II. In untransfected cells, stimulation with IFN-I or -II induced STAT1 redistribution to the nucleus (**Figure 2A**), whereas only IFN-I did so for STAT2 (**Figure 2B**). In contrast, translocation of STAT1 and 2 was blocked in cells expressing 018 (**Figure 2A, B)**. Consistent with previous reports, NiV-V also blocked STAT1 translocation, but unlike 018, NiV-V redistributed STAT1 to a predominantly cytoplasmic localisation in non-stimulated cells due to harbouring a nuclear export signal (Rodriguez et al., 2004) (**Figure 2A**).

**Figure 2.**
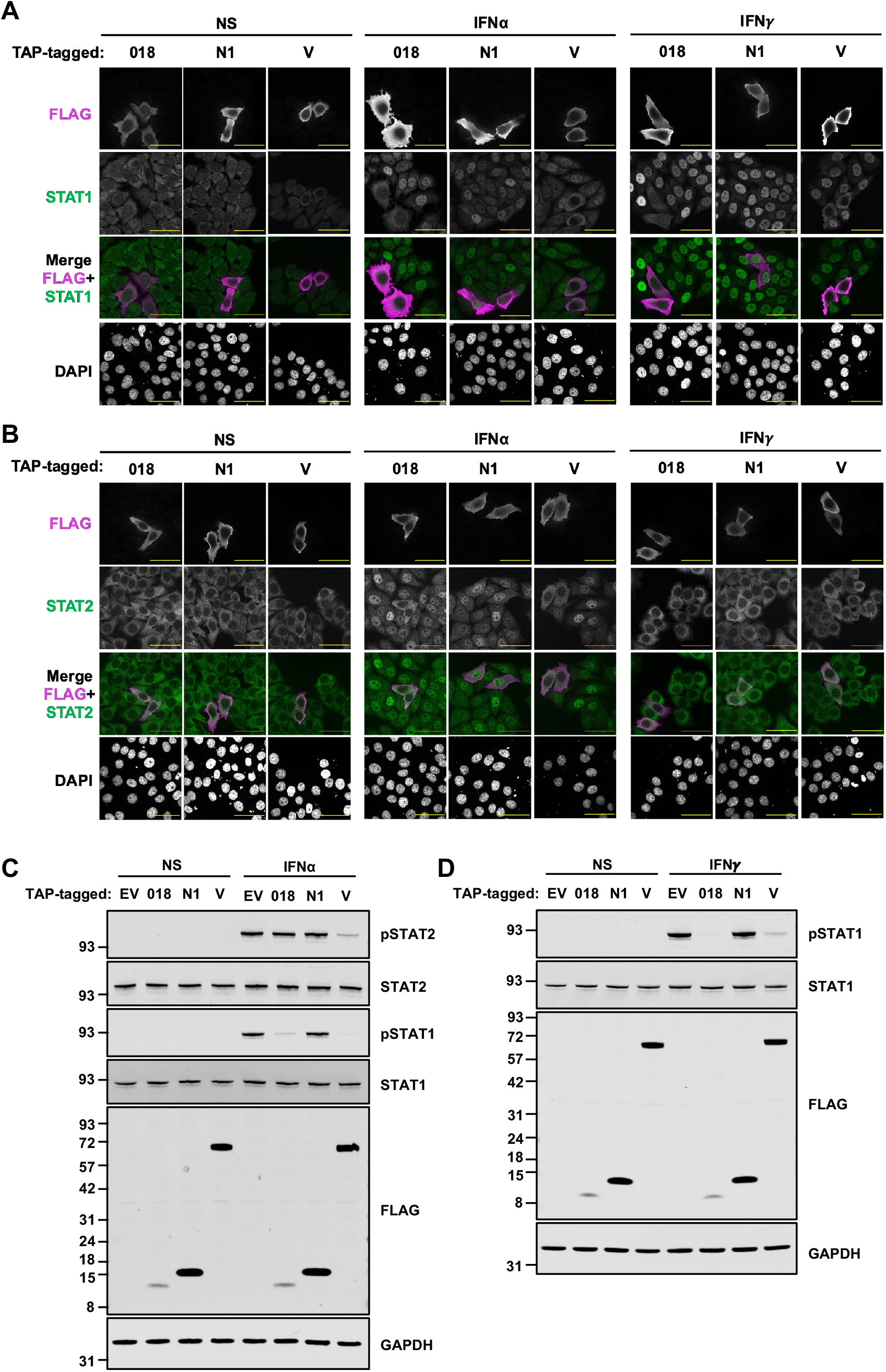
Phosphorylation of STAT1 at Tyr 701 is blocked by 018. (**A-B**) HeLa cells were transfected with plasmids expressing TAP-tagged 018, N1 or NiV-V and then were stimulated with IFNα (1000 U/mL) or IFNγ (25 ng/mL) for 1 h. Cells were fixed and permeabilised, and then immunostained with α-FLAG (pink) (**A-B**) and either α-STAT1 (green) (**A**) or α-STAT2 (green) (**B**) and mounted in Mowiol containing DAPI to stain DNA. Cells were visualised by confocal microscopy. Scale bar (yellow) = 50 µm. (**C-D**) T-REx 293 cells were induced with dox (100 ng/mL) to express indicated proteins. Cells were stimulated with IFNα (1000 U/mL) (**C**) or IFNγ (25 ng/mL) (**D**) for 30 min and lysates were analysed by immunoblotting against proteins/epitope indicated. Quantification of band intensities for (**C-D**) is provided in **Figure S2**. Data for (**A-B**) and (**C-D**) are representative of two or three individual experiments, respectively.

Prior to translocation, STATs are phosphorylated at conserved tyrosine residues near their C terminus. To test if 018 inhibited phosphorylation of STAT1 at Tyr701 (pSTAT1) and STAT2 at Tyr-690 (pSTAT2), T-REx 293 cells expressing TAP-tagged 018 or controls were stimulated with IFN-I or -II and lysates were analysed by immunoblotting. IFN-I stimulation increased pSTAT1 and pSTAT2 levels in both EV and N1-expressing cells, whereas cells expressing 018 showed very low levels of pSTAT1 (**Figure 2C, Figure S2A**). In contrast, 018 only affected pSTAT2 levels marginally (**Figure 2C, Figure S2B**). Consistent with previous reports, NiV-V blocked STAT1 phosphorylation (Rodriguez et al., 2002). Interestingly, NiV-V also blocked STAT2 phosphorylation, which has not been reported previously, but is consistent with NiV-V harbouring a distinct STAT2-binding site (Rodriguez et al., 2004). IFN-II increased pSTAT1 in control cells, whereas STAT1 phosphorylation was blocked by 018 (and NiV-V) (**Figure 2D, Figure S2C**). These data show 018 blocks phosphorylation of STAT1 at Tyr701 after stimulation with IFN-I or -II and thus prevents STAT1/2 translocation.

### A minimal 21 aa fragment of 018 is sufficient to bind STAT1

Next, we assessed if 018 interacts with cellular proteins involved in IFN signal transduction. TAP-tagged 018 was expressed in 2fTGH cells and its ability to co-precipitate STAT1, STAT2 and IRF9 was assessed by pulldown. 018 co-precipitated STAT1 and to a lesser degree STAT2, whereas no interaction with IRF9 was observed (**Figure 3A**). In 2fTGH-derived STAT1 knockout cells (U3A), the 018:STAT2 interaction was lost, indicating the interaction was either indirect via STAT1, or STAT1 was required for 018 to precipitate STAT2 (**Figure 3B**). In 2fTGH-derived STAT2 knockout cells (U6A), the 018:STAT1 interaction was retained (**Figure 3B**). The 018:STAT1 interaction was confirmed to be direct as 018 and STAT1 still co-precipitated when expressed by a cell-free based transcription and translation system (**Figure S3A**).

**Figure 3.**
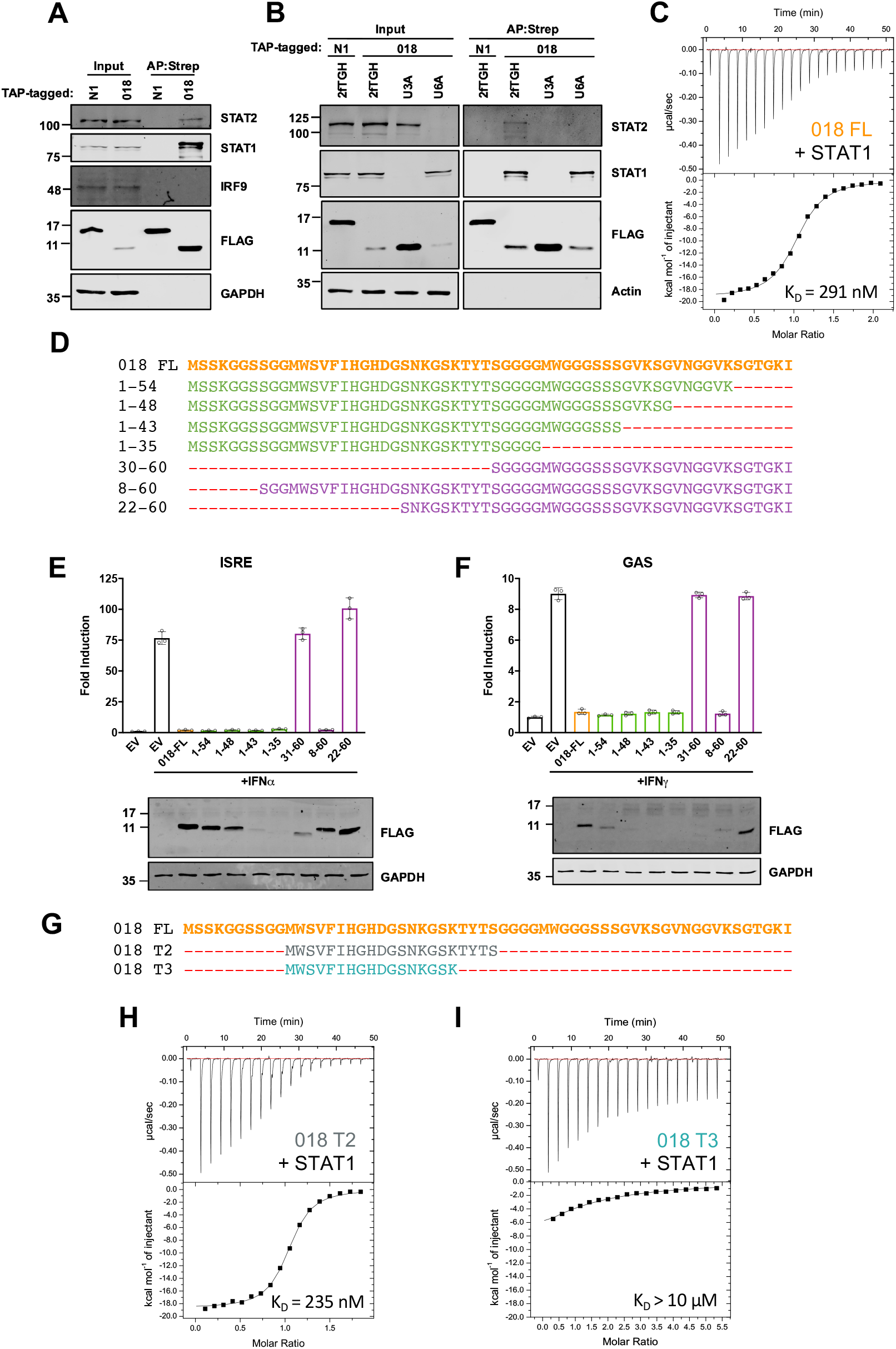
A minimal 21 aa fragment of 018 is sufficient to bind STAT1. (**A-B**) TAP-tagged 018 and N1 were expressed in 2fTGH cells (**A**) or 2fTGH, U3A (STAT1^-/-^ ) and U6A (STAT2^-/-^) cells (**B**) by transfection and were affinity purified by Strep-Tactin. Whole cell lysate (Input) and affinity purified proteins (AP:Strep) were analysed by immunoblotting. **(C)** ITC data for GB1-018 (100 µM) titrated into U-STAT1 (10 µM). Fitting of the isotherm (bottom) to a one site model gave a KD of 290 nM. Initial low volume injection is excluded from analysis. Complete fitted ITC parameters are provided in **Table S5**. (**D**) Sequences for TAP-tagged C-terminal (green) and N-terminal (purple) 018 truncation mutants. **(E)** HEK 293T or **(F)** HeLa cells were transfected with reporter plasmids ISRE-Luc **(E)** or GAS-Luc **(F)** along with *TK*-*Renilla* and vectors from (**D**). Cells were stimulated with IFNα (1000 U/mL) (**E**) or IFNγ (25 ng/mL) (**F**) for 6 (**E**) or 8 h (**F**) and then luciferase activity was measured. Means ± SD (n=3 per condition) are shown. Immunoblots were stained with antibodies against proteins/epitopes indicated (**A-B**) and (**E-F**). Percentage inhibitory activity and relative protein expression levels from figures (**E-F)** are provided in **Figure S3**. Data shown in (**A-B**) and (**E-F**) are representative of three or two individual experiments, respectively. (**G**) Sequences for purified GB1-fused 018 truncation mutants. (**H-I**) ITC data for 150 µM GB1-018^T2^ (**H**) or 350 µM GB1-018^T3^ (**I**) titrated into 150 µM STAT1. Accurate fitting of the isotherm for (**I**) was not possible due to the low C-value of the reaction.

To characterise the 018:STAT1 interaction, each protein was expressed and purified from *E. coli*. 018 was fused to the B1 domain of protein G (GB1) to improve solubility and expression. Using isothermal titration calorimetry (ITC), we observed a KD value of 291 nM, with a stoichiometry of 1.02, indicating a single 018 molecule binds per U-STAT1 protomer (**Figure 3C**). The effect of 018 on U-STAT1 quaternary assembly was evaluated by SEC-MALS. U- STAT1 alone eluted predominantly as tetrameric and dimeric species, and preincubation with excess GB1-018 resulted in the two peaks having earlier elution volumes and increased masses (**Figure S3B**). This indicates that 018 binds U-STAT1 without altering its oligomeric state.

Next, the region of 018 required to inhibit IFN signalling was mapped. C-terminal and N-terminal truncations of 018 were constructed and their ability to inhibit IFN-I and -II signalling was assessed (**Figure 3D-F**). For ease of interpretation, inhibitory activity was categorised as (i) greater than 95%, (ii) between 75% and 95%, or (iii) less than 25%, which was deemed to be non-inhibitory. Mutant 1-35 had the largest C-terminal truncation and still demonstrated >95% inhibition (**Figure 3E,F**). Mutant 8-60 inhibited >95%, whereas mutant 22-60 lost inhibitory activity (<25%) and showed comparable expression to the full-length protein (**Figure 3E,F**). These data show that aa 8-35 of 018 is sufficient for pathway inhibition.

To refine the region of 018 responsible for pathway inhibition, additional mutants, truncating inwards from aa 8 and 35 were constructed (**Figure S3C**). Mutant 11-60 retained >95% inhibition whereas mutants with further N-terminal truncation had reduced inhibitory activity despite WT expression levels (**Figure S3D,E**). Mutant 1-30 inhibited between 75%-95%, demonstrating a marginal loss in inhibitory activity, however, expression was undetectable (**Figure S3D,E**). All further C-terminal truncations showed <25% inhibitory activity, but again, expression was undetectable (**Figure S3D,E**). The same pattern of inhibitory activity by 018 truncations mutants was observed for both IFN-I and -II signalling (**Figure S3F**), indicating the same region of 018 is required to inhibit both pathways.

These observations map a putative minimal inhibitory region of 018 to aa 11-31. The C-terminal boundary was defined assuming the slight reduction in inhibitory activity after deletion of residues 35-31 was due to lower protein expression levels, whereas further truncation removed functional residues. Ser31 is included within the minimal inhibitory region as it is highly conserved in orthopoxvirus orthologues of 018 (**Figure S1)**.

ITC measurements of the putative minimal fragment (018^T2^) with STAT1 gave a KD of 235 nM, a value comparable to that of full length 018 (291 nM) (**Figure 3H)**. Removal of the C-terminal 28-TYTS-31 (018^T3^) from the putative minimal fragment led to a large reduction in affinity (>10 μM), thereby demonstrating the importance of these residues (**Figure 3I)**. Collectively, these data show that a 21-residue segment of 018, aa 11-31, is sufficient for maximal STAT1 binding and inhibitory activity.

### 018 is a virulence factor

To study the role of 018 during infection, a VACV (strain WR) 018 deletion mutant (termed vΔ018) was constructed. The wild-type sibling virus (termed v018) and vΔ018 were analysed by PCR (**Figure S4A**) and genomic sequencing, which showed no differences other than the intended deletion of the 018 ORF. Comparison of v018 and vΔ018 in cell culture displayed no difference in replication or plaque size (**Figure S4B-D**). Another VACV (termed vTAP-018) was constructed by reintroduction of the 018 ORF fused to an N-terminal TAP-tag into vΔ018 at its natural locus. Pulldown of TAP-tagged 018 expressed from vTAP-018 confirmed the 018:STAT1 interaction during infection (**Figure S4E, F**).

Next, vΔ018’s ability to inhibit IFN signalling was assessed. A549 cells were infected with v018 or vΔ018 and, at the indicated times p.i., were stimulated with either IFN-I or -II, after which the pSTAT1 level was determined by immunoblotting. Cells were washed once prior to stimulation to remove the majority of soluble VACV IFN decoy receptors B8 and B18. This, however, will not fully remove B18 (IFN-I decoy receptor) due to its ability to bind to the cell surface (Alcamí et al., 2000). Although by 2 h p.i., both v018 and vΔ018 inhibited pSTAT1 induction after IFN-I stimulation, v018 inhibited earlier and to a greater extent. **(Figure 4A)**. In contrast, pSTAT1 induction was inhibited by v018 but almost fully rescued to mock levels in vΔ018-infected cells after IFN-II stimulation **(Figure 4B)**. Consistent with this finding, STAT1 translocation was blocked by v018 after IFN−ΙΙ stimulation, whereas in vΔ018-infected cells, STAT1 was predominantly nuclear **(Figure 4C)**. The impaired ability of vΔ018 to inhibit IFN-II signalling was illustrated further by increased IRF1 levels (a canonical IFNγ ISG) in cells infected with v018 compared to vΔ018 after IFN-II stimulation at both the mRNA (**Figure S4G**) and protein level (**Figure 4D**).

**Figure 4.**
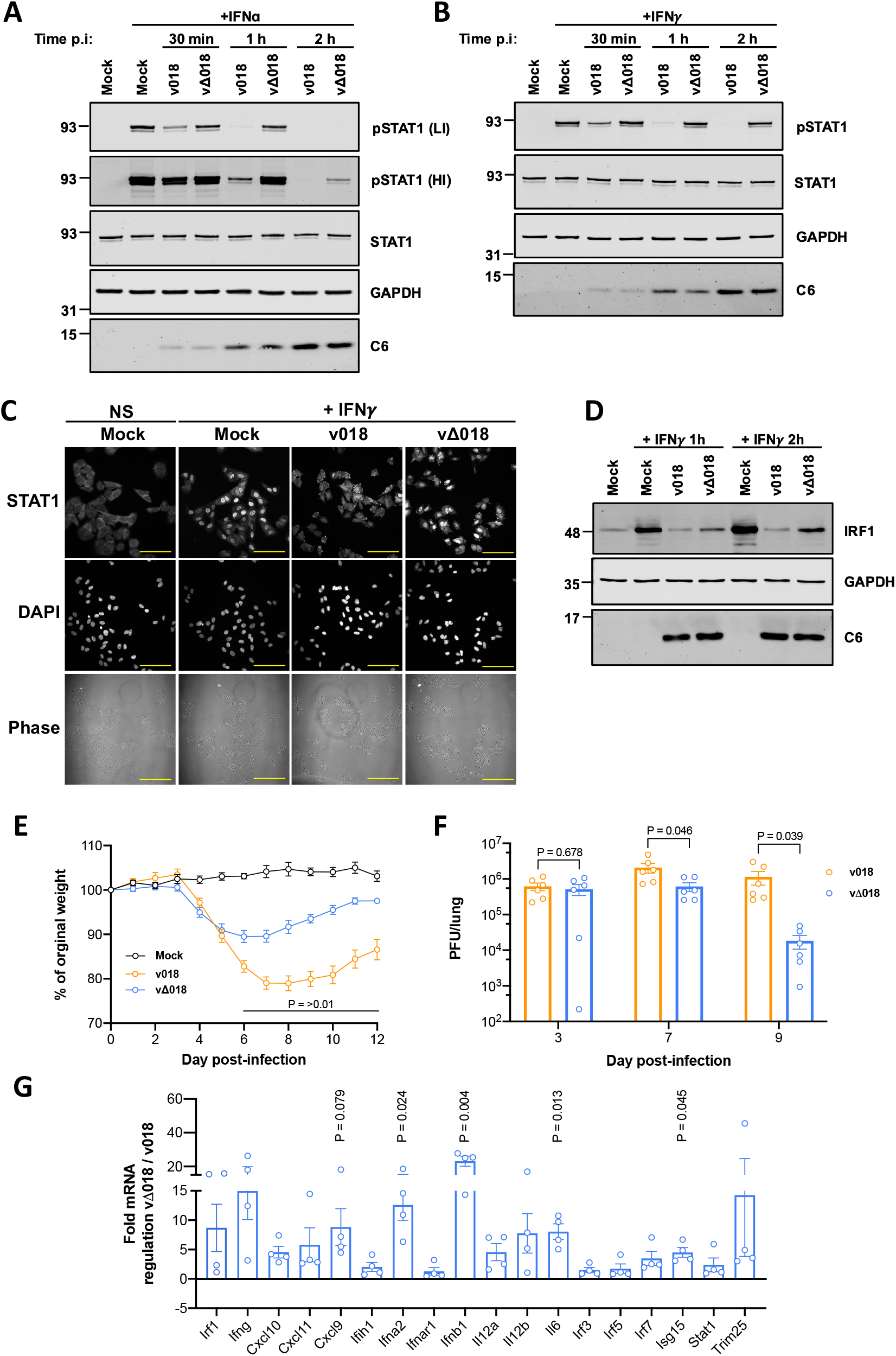
018 is a virulence factor. (**A-B**) A549 cells were mock infected or infected with v018 or vΔ018 at 10 pfu/cell. At 30 min, 1 h or 2 h post infection (p.i.) cells were washed once, then stimulated with IFNα (1000 U/mL) (**A**) or IFNγ (25 ng/mL) (**B**) for 30 mins and lysates were analysed by immunoblotting. (**C-D**) A549 cells were infected as described for (**A-B**) and at 2 h p.i., cells were washed once, and then stimulated with IFNγ (25 ng/mL) for 30 mins (**C**), or 1 and 2 h (**D**). (**C**) Cells were fixed and permeabilised then immunostained with α-STAT1 and mounted in Mowiol-containing DAPI to stain DNA and visualised by confocal microscopy. Scale bar (yellow) = 100 µm. (**D**) Cell lysates were analysed by immunoblotting. Immunoblots were stained against proteins indicated including the early VACV protein C6 to control for equal infection (**A-B** and **D**). For (**A**) high intensity (HI) and low intensity (LI) scans for α-pSTAT1 are shown. Data for (**A-D**) are representative of three individual experiments. (**E-G**) Female BALB/c mice 6-10 weeks old were infected via the intranasal route with either v018 (orange) or vΔ018 (blue) at 10^3^ (**E- F**) or 10^5^ (**G**) pfu and weight was measured daily (**E**) or virus titres of upper lungs lobes were measured by plaque assay on days 3, 7 and 9 (**F**), or mice were sacrificed at 3 day p.i. and mRNA levels of indicated genes isolated from upper lung lobes were analysed by RT-qPCR (**G**). Data from (**E-F**) are representative of at two individual experiments using either 5 or 3 mice, respectively, per group which were then pooled. Data from (**G**) is representative of 4 (vΔ018) or 3 (v018) mice per group. For (**E-G**) means ± SEM are shown, and P values were calculated using Unpaired T-test with (**E-F**) or without (**G**) Welch’s correction.

To evaluate if 018 contributes to virulence, BALB/c mice were infected via the intranasal route with either v018 or vΔ018 and their weight was measured daily (**Figure 4E)**. Mice infected with vΔ018 lost significantly less weight than those infected with v018 (**Figure 4E)** and showed reduced virus titres at 7 and 9 days p.i. (**Figure 4F**). Furthermore, consistent with 018 functioning as an immunomodulator, mRNAs for several ISGs, chemokines and IFNs were upregulated in the lungs of mice infected with vΔ018 compared to v018 (**Figure 4G)**. Collectively, these data show VACV lacking 018 is defective in inhibition of IFN-induced signalling and is attenuated in mice.

### 018 binds the STAT1 SH2 domain to block its association with the phosphorylated IFNGR1

To identify which STAT1 domain/s 018 binds, the ability of 018 to interact with several STAT1 truncations and STAT1-STAT3 chimeras was tested (**Figure 5A**). 018 bound a chimera with linker domain (LD), SH2 and transactivation domain (TAD) of STAT1 (31F), but not a chimera with coiled-coil and DNA-binding domains of STAT1 (13F) (**Figure 5B**). These chimeras have been studied with NiV-V, which also only binds 31F (Rodriguez et al., 2004). 018 bound STAT1 C-terminal truncations that lack the final 38 residues (STAT1β, a naturally occurring isoform of STAT1) or the entire TAD (**Figure 5C**). Lastly, 018 bound a chimera that contained only the SH2 and TAD of STAT1 (Fus 1) but not a chimera that contained the LD of STAT1 (Fus 2) nor with STAT3 alone (**Figure 5D**). Together, these pulldowns show 018 binds the SH2 domain of STAT1.

**Figure 5.**
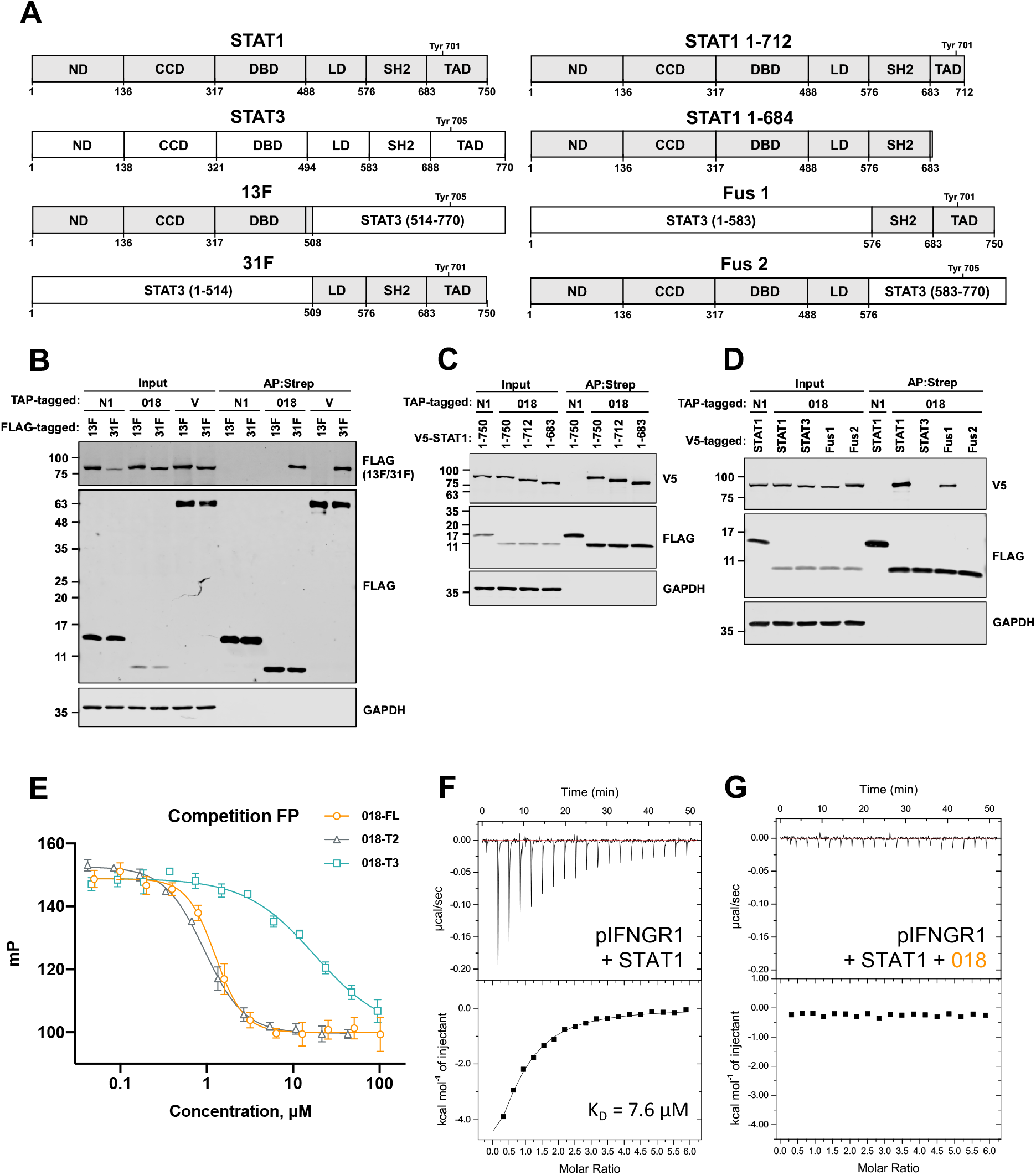
018 binds the STAT1 SH2 domain to block its association with the phosphorylated IFNGR1. (**A**) Schematic of STAT1-STAT3 chimeras and STAT1 truncation mutants. STAT1 regions (grey) and STAT3 (white) are shown, and domains are annotated as ND (N-terminal domain), CCD (Coiled-coil domain), DBD (DNA-binding domain), LD (Linker-domain), SH2 (Src homology-2-domain), and TAD (Transactivation-domain). (**B-D**) TAP-tagged proteins indicated were co-expressed with either FLAG (**B**) or V5-tagged (**C-D**) STAT proteins from (**A**) by transfection in U3A (STAT1^-/-^) cells and TAP-tagged proteins were purified by Strep- Tactin. Whole cell lysates (Input) and affinity-purified proteins (AP:Strep) were analysed by immunoblotting for the indicated proteins/epitopes. Data from (**B-D**) are representative of two individual experiments. (**E**) Competition FP measurements for GB1-018 and truncation mutants. Each reaction contained 10 nM fluorescein-pIFNGR1 12-mer preincubated with 1.5 µM U-STAT1, to which two-fold serial dilutions of purified GB1-018 proteins were added (all concentrations are final values). One hundred mP represents the calibrated FP value of the free fluorescent probe. (**F-G**) ITC data for 300 µM pIFNGR1 5-mer titrated into 100µM U-STAT1 (**F**) or 100µM U-STAT1 preincubated with GB1-018 (**G**). No heat of binding was detected for (**G**). Complete fitted ITC parameters are provided in **Table S5**.

The finding that 018 binds the SH2 domain allowed us to hypothesise how 018 blocks STAT1 phosphorylation. Given that 018 inhibition of IFN-II signalling during infection was non-redundant, we focused on this pathway to study 018 mechanistically. The IFNGR is composed of two ligand-binding IFNGR1 chains and two IFNGR2 chains and binds dimeric IFNγ (Mendoza et al., 2019). Ligand engagement induces JAK-1 phosphorylation of IFNGR1 at Tyr440 (Briscoe et al., 1996; Greenlund et al., 1994). STAT1 then docks at the IFNGR1 pTyr site via its SH2 domain and is itself phosphorylated (Greenlund et al., 1995). pSTAT1 then undergoes structural rearrangement to a parallel dimer orientation and dissociates from the receptor. We hypothesised that by binding the SH2 domain, 018 blocks STAT1 recruitment to pIFNGR1 and thus prevents STAT1 phosphorylation.

To test this, a fluorescence polarisation (FP) assay was established using a fluorescent 12-mer peptide corresponding to the pIFNGR1 sequence harbouring the STAT1 docking site (pYDKPH) as a probe. Addition of 018 to a preformed STAT1-pIFNGR1 probe led to a dose-dependent displacement of probe and an IC50 value of 1.26 μM (**Figure 5E**). IC50 values of 0.93 μM and 17.82 μM for 018^T2^ and 018^T3^, respectively, were obtained, demonstrating that 018^T2^, but not 018^T3^, has comparable inhibitory activity to full length 018, consistent with ITC data (**Figure 5E**).

The mechanism was further validated by competition ITC. A 5-mer peptide corresponding to the pIFNGR1 docking region (pYDKPH) was titrated into U-STAT1, giving a KD value of 7.6 μM (**Figure 5F**). In contrast, inclusion of excess 018 in the calorimeter cell resulted in complete loss of detectable binding (**Figure 5G**). Taken together, these data demonstrate a competitive inhibition mechanism, whereby 018 binds the SH2 of STAT1 and prevents STAT1 from engaging the active IFN signalling receptor complex.

### Vaccinia 018 and Nipah virus V protein utilise a shared motif to engage STAT1

NiV-V, W and P proteins, encoded by the P gene, all inhibit IFN signalling. They have distinct C-terminal sequences but share a common 407 aa N-terminal region to which the IFN inhibitory activity was mapped (aa 114-140) (Ciancanelli et al., 2009). Here, we focus on this STAT1-binding region and refer to this within NiV-V.

The observation that 018 and NiV-V bind STAT1, block STAT1 phosphorylation and bind the 31F chimera suggested they might share a similar mode of action. Alignment of the NiV-V STAT1 binding region and 018^T2^ revealed aa similarity exemplified by a conserved HxH motif preceded by a cluster of conserved hydrophobic residues (**Figure 6A**). Recent ITC data showed a NiV-V fragment (aa 92-190) binds STAT1 directly but weakly (KD >100 μM) and mutation of 117-HDH-119 to 117-AAA-119 abolished binding (Jensen et al., 2020). To determine whether the HxH motif of 018 had analogous function, we mutated 17-HGH-19 to 17-AGA- 19 (018^AGA^). Unlike 018, 018^AGA^ did not co-precipitate with endogenous STAT1 in cells (**Figure S5**). Furthermore, ITC titration of purified GB1-fused 018^AGA^ into STAT1 resulted in no detectable binding (**Figure 6B**). Loss in STAT1 binding ability correlated with a loss of inhibitory activity because 018^AGA^ was unable to inhibit IFN-I and -II signalling by reporter gene assay (**Figure 6C,D**). Consistent with this, 018^AGA^ did not interfere with STAT1:pIFNGR1 12-mer interaction by FP (**Figure 6G**). In addition, 018^AGA^ showed no inhibition of STAT1-pIFNGR1 binding via ITC (**Figure 6H**).

**Figure 6.**
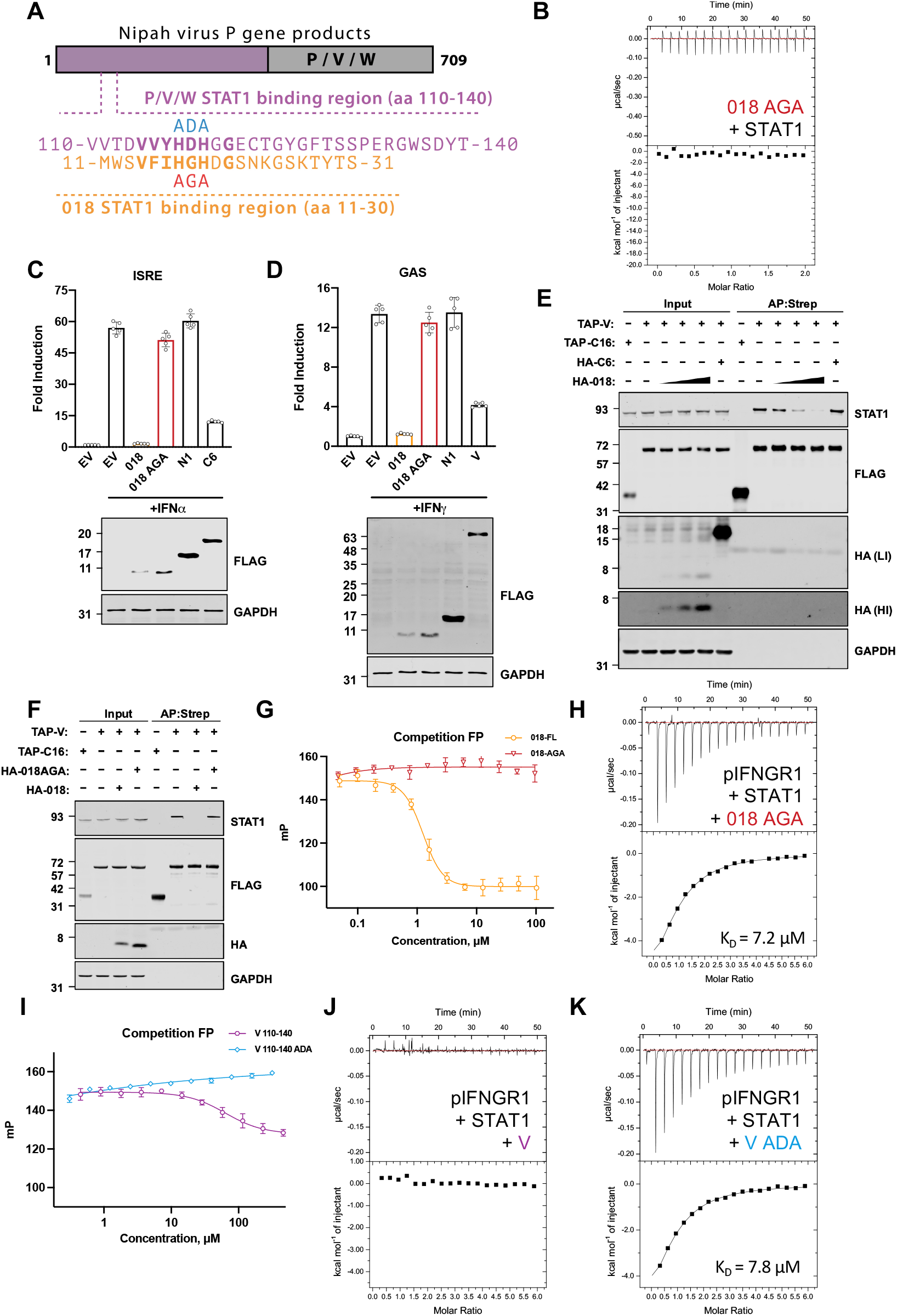
Vaccinia 018 and Nipah virus V protein utilise a shared motif to engage STAT1. **(A)** Schematic of Nipah virus P, V and W proteins encoded by the P gene indicating the common N-terminal region (purple) and unique C-terminal region (dark grey). Below, the STAT1-binding regions of P/V/W (residues 110-140, purple) and 018 (residues. 11-31, orange) are aligned with key conserved residues highlighted in bold. Sites of NiV-V^ADA^ (blue) and 018^AGA^ (red) mutants are shown. (**B**) ITC data for the titration of 100 µM GB1-018^AGA^ into 10 µM U-STAT1. No heat of binding was observed. (**C**) HEK 293T or (**D**) HeLa cells were transfected with reporter plasmids ISRE-Luc **(C)** or GAS-Luc (**D**) along with *TK-Renilla* and vectors expressing the proteins indicated fused to a TAP-tag. Cells were stimulated with IFNα (1000 U/mL) **(C)**, or IFNγ (25 ng/mL) **(D)** for 6 (**C**) or 8 h (**D**) and then luciferase activity was measured. Means ± SD (n=5 per condition) are shown. (**E-F**) TAP-tagged and HA-tagged proteins were co-expressed in HEK 293T cells by transfection as indicted and TAP-tagged proteins were affinity purified by Strep-Tactin. Whole cell lysates (Input) and affinity purified (AP:Strep) proteins were analysed by immunoblotting. For (**E**) high-intensity (HI) and low intensity (LI) scans are shown for α-HA. VACV proteins TAP-C16 and HA-C6 were used as a pulldown and competition protein controls respectively. Immunoblots were stained against proteins/epitopes indicated (**C-F**). Data shown in (**C-D**) and (**E-F**) are representative of two or three individual experiments respectively. (**G**) Competition FP measurements for GB1-018 and GB1-018^AGA^ binding to U-STAT1. Each reaction contained 10 nM fluorescein-pIFNGR1 12- mer preincubated with 1.5 µM U-STAT1, to which two-fold serial dilutions of purified GB1- 018 proteins were added (all concentrations are final values). One hundred mP represents the calibrated FP value of the free fluorescent probe. (**H+J+K**) ITC data for 300 µM pIFNGR1 5- mer titrated into 10 µM U-STAT1 preincubated with 50 µM GB1-018^AGA^ (**H**), 200 µM NiV- V (**J**) or 200 µM NiV-V^ADA^ (**K**). No heat of binding was detected for the reaction containing GB1-NiV-V. Complete fitted ITC parameters are provided in **Table S5**. (**I**) Competition FP measurements for GB1-NiV-V and GB1-NiV-V^ADA^ binding to U-STAT1. Each reaction contained 10 nM fluorescein-pIFNGR1 12-mer preincubated with 1.5 µM U-STAT1, to which two-fold serial dilutions of purified GB1-NiV constructs were added (all concentrations are final values). The NiV-V^ADA^ curve has a positive slope at high protein concentrations due to either increased sample viscosity or non-specific interactions. One hundred mP represents the calibrated FP value of free fluorescent probe.

Consistent with the idea that 018 and NiV-V harbour analogous motifs, recently the site for NiV-V binding to STAT1 was mapped to the SH2 domain of STAT1 (Keiffer et al., 2020). To assess if these viral proteins target the same SH2 interface, the ability of 018 to outcompete the NiV-V:STAT1 interaction was tested. In cells transfected with TAP-tagged NiV-V, NiV-V co-precipitated with endogenous STAT1, however, this was decreased in a dose-dependent manner by expression of HA-tagged 018 (**Figure 6E**). In contrast, HA-tagged 018^AGA^ did not affect the NiV-V:STAT1 interaction (**Figure 6F**). These data show that 018 and NiV-V utilise a shared motif to bind a common interface on the SH2 domain of STAT1.

Previous reports show NiV-V sequesters STAT1 and 2 within the cytoplasm and prevents STAT1 phosphorylation (Rodriguez et al., 2002). The finding that 018 and NiV-V bind STAT1 via the same interface prompted us to assess if, like 018, NiV-V competes with pIFNGR1 to bind STAT1. To test this, NiV-V STAT1-binding fragment residues 110-140 (NiV-V^110-140^) fused to a GB1 tag was purified together with a mutant in which His117 and His119 of the HxH motif were mutated to Ala (NiV-V^ADA^). By FP assay, addition of NiV-V to the preformed STAT1-pIFNGR1 12-mer complex led to a modest reduction in polarisation, whereas addition of NiV-V^ADA^ was non-competitive (**Figure 6I**). Consistent with these data, preincubation of STAT1 with NiV-V abolished any detectable binding between STAT1 and the pIFNGR 5-mer by ITC (**Figure 6J**). In contrast, preincubation with NiV-V^ADA^ did not prevent STAT1:pIFNGR binding (**Figure 6K**). These data show that, in the context of IFN-II signalling, NiV-V can block STAT1 recruitment to the active IFNGR signalling complex.

### Phosphotyrosine pocket-independent binding of 018 to the STAT1 SH2 domain

A feature of the SH2 interface is a deep pTyr pocket that binds the phosphate group and the phenyl ring of phosphotyrosine. Remarkably, 018 binds the STAT1 SH2 domain with high-affinity and competes with pIFNGR1 without a pTyr modification. Intrigued by this observation, we crystallised the STAT1 132-684 core fragment complexed with the minimal 018 peptide (Met11-Ser31). Crystals diffracted to 2.0 Å with 018 electron density clearly defined for most of the peptide, with only Ser31 not visible (**Figure S6A**).

The 018 peptide forms a β-hairpin fold with a β-turn midway through the sequence (**Figure 7A,B**) and the two strands of the peptide augment the central β-sheet of the SH2 domain, with Val14-His17 backbone hydrogen-bonding to the βD strand of the SH2 domain (**Figure 7C**). There is spatial overlap with published binding modes of pTyr peptides from pIFNGR1 and pSTAT1 homodimer (**Figure 7B**). The 680 Å^2^ interface is formed by a large number of shallow contacts exclusively within the SH2 domain. Residues Trp12, Val14, Ile16 comprise a continuous hydrophobic interface with STAT1 helix αA and strand βD (**Figure 7D**). This is followed by a HxH motif, in which His17 forms an imidazole-to-imidazole hydrogen bond with His629 of STAT1 (**Figure 7D, E**). The His17 rotamer is stabilised intramolecularly by a second hydrogen bond with the backbone carbonyl of 018 Gly21. Gly18 carbonyl forms a hydrogen bond with Tyr651 hydroxyl of STAT1, similar to pIFNGR1 Pro443 (PDB: 1YVL). His19 occupies the same cleft as His444 of pIFNGR1, forming an identical π-stacking interaction with STAT1 Tyr634. The Asp20 sidechain stabilises the β-turn by hydrogen bonding with the Ser21 backbone and forms an intramolecular salt bridge with Lys24 (**Figure 7D**). An inter-strand hydrogen bond between the hydroxyl groups of Ser13 and Thr28 act as a non-covalent bridge that may stabilise the β-hairpin fold (**Figure 7D**).

**Figure 7.**
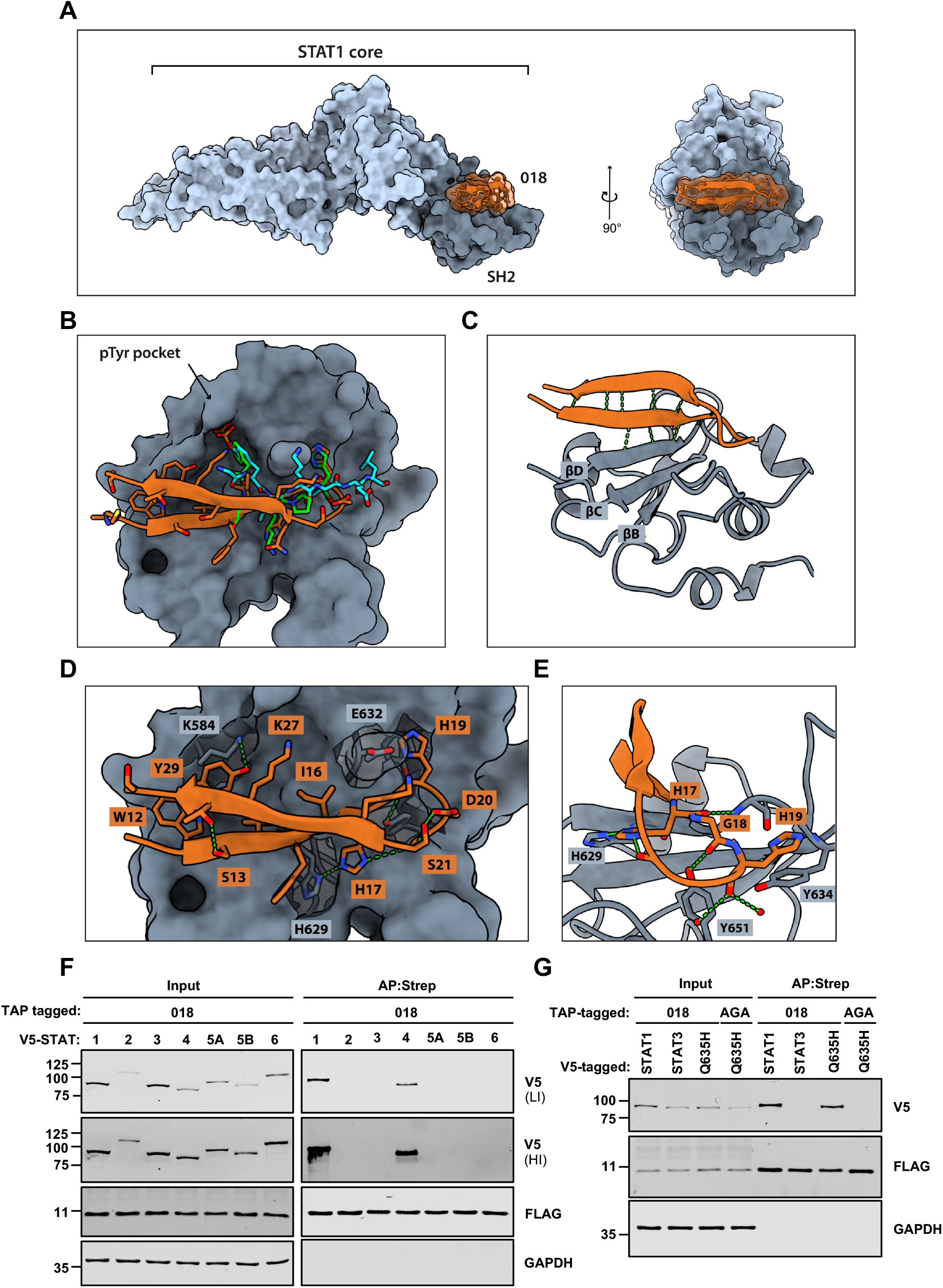
Structural basis of 018 binding to U-STAT1. A crystal structure was determined for 018:STAT1 core fragment complex (PDB: 7nuf). In all images, 018 is depicted in orange, the SH2 domain is dark grey and the rest of the core fragment is light grey. (**A**) Surface representation of the complex viewed from two perpendicular axes. 018 binding mode at the STAT1 SH2 domain superimposed with IFNGR1 phosphopeptide (green, PDB: 1yvl) and STAT1 pTyr701 phosphopeptide (cyan, PDB: 1bf5). (**C**) Ribbon representation of 018 and the STAT1 SH2 domain with β-sheet-forming hydrogen bonds depicted in green. SH2 domain core β-strands are labelled according to conventional nomenclature. (**D**) Detailed depiction of 018 binding to the STAT1 SH2 domain. 018 sidechains are depicted as sticks, while backbone atoms are represented by a ribbon cartoon. Key STAT1 sidechains are depicted as sticks under semi-transparent surface. (**E)** A zoomed view of HxH motif binding. (**F, G**) V5-tagged and TAP-tagged proteins were co-expressed in U3A (STAT1^-/-^) cells by transfection as indicted in the figure and TAP-tagged proteins were affinity purified using Strep-Tactin. Whole cell lysates (Input) and affinity purified proteins (AP:Strep) were analysed by immunoblotting against the indicated proteins/epitope. STAT3^Q635H^ and 018^AGA^ are labelled as Q635H and AGA, respectively (**G**). For (**F**) high (HI) and low intensity (LI) scans of α-V5 are shown. Data shown in (**F, G**) are representative of two individual experiments.

Strikingly, 018 does not interact with the pTyr pocket. The only tyrosine in the peptide, Tyr29, hydrogen bonds with the ζ-amine of STAT1 Lys584 through its hydroxyl and makes van der Waals contacts with the alkyl chain of the same lysine (**Figure 7D**). The lower affinity of 018^T3^ compared to 018^T2^ may result from the loss of interactions made by Thr28 and Tyr29.

### A single histidine found in STAT1 and 4 is a key determinant of 018 selectivity

High sequence similarity between SH2 domains of STATs led us to investigate if 018 interacts with other STATs. In humans, there are seven STATs (STAT1, 2, 3, 4, 5A, 5B and 6) (Ihle, 2001). U3A cells were co-transfected with V5-tagged STAT1-6 and TAP-tagged 018. Pulldown of 018 demonstrated that 018 binds STAT1 and STAT4, but not other STATs (**Figure 7F**).

To understand the observed specificity of 018 for STAT1 and 4, STAT SH2 domain alignments were integrated with our structural data (**Figure S6B**). In the crystal structure, a specific interaction between 018 His17 and His629 of STAT1 was observed. Only STAT1 and 4 have a histidine at this position, and so other STATs would fail to recapitulate this interaction. To test if STAT1 His629 was critical for specificity, a STAT3 mutant was made in which the structurally equivalent Glu635 was mutated to His. This enabled 018 to co-precipitate STAT3^Q635H^, allowing the unambiguous assignment of specificity determinants of 018 binding (**Figure 7G**).

## Discussion

STAT1 and 2 are central to IFN signalling and thus are common targets for viral antagonism (Harrison and Moseley, 2020), however structural details of STAT:antagonist complexes have remained elusive with a few exceptions. The complex of SeV C protein with the N-terminal domain of STAT1 indicates that C protein interferes with the oligomeric state of STAT1 (Oda et al., 2015) whilst the structures of dengue and Zika virus NS5 proteins in complex with STAT2 revealed that both NS5 proteins overlap the IRF9 binding site to prevent ISGF3 assembly (Wang et al., 2020). A similar mechanism was described for measles V protein (Nagano et al., 2020). Here, the structure of 018, an uncharacterised poxvirus protein, with STAT1, shows 018 occupies the STAT1 SH2 domain to block STAT1 association with the active pIFNGR.

The molecular recognition between STAT SH2 domains and a pTyr site represents a conserved mechanism for STAT recruitment to activated receptors. The pTyr is estimated to contribute half the binding energy, while the remaining specificity is provided by a small number of adjacent residues (Kaneko et al., 2010; Ladbury and Arold, 2011). For STAT1, 018 overlaps with these specificity-determining sites and obstructs the pTyr pocket without occupying it.

To establish if a similar binding mode exists, we examined 524 SH2-containing structures retrieved from PDB based on either Pfam or SMART annotation. The majority of liganded SH2 domains bind a pTyr-containing peptide, a synthetic pTyr mimetic or an unphosphorylated tyrosine at the pTyr pocket. Several structures contain SH2 domains as part of a larger protein-protein interaction, in which the pTyr pocket is not occupied, however, in such cases, the interface extends significantly beyond the SH2 phosphopeptide site. The closest binding mode analogue to 018 was a monobody that binds at the phosphopeptide site of SHP-1 phosphatase without interacting with the pTyr pocket itself (PDB: 6SM5). Hence, we suggest 018 has an unprecedented mode of high-affinity SH2 domain binding.

For IFN-I-induced signalling, 018 blocked pSTAT1 induction but only modulated pSTAT2 levels minimally. After IFN-I stimulation, STAT2 docks at pTyr466 on IFNAR1 and subsequently is phosphorylated at Tyr690 (Yan et al., 1996). The pTyr690 of STAT2 serves as a docking site for STAT1 to present STAT1 for proximal phosphorylation at Tyr701 by JAKs (Leung et al., 1995; Li et al., 1997; Qureshi et al., 1996). Additional Tyr phosphorylation sites on IFNAR2 are also important for ISGF3 formation and could serve as docking sites for STAT1 and 2 functioning in a cell type- or species-dependent manner (Zhao et al., 2008). Thus, we rationalise that during IFN-I signalling, occupancy of the STAT1 SH2 domain by 018 would diminish STAT1 engagement of either STAT2 pTyr690 or IFNAR to prevent STAT1 phosphorylation.

STAT4 was identified as an additional binding partner of 018. STAT4 is activated by phosphorylation predominantly in response to IL-12 and IFN-I and promotes IFNγ production during virus infection (Nguyen et al., 2002; Yang et al., 2020). The activation of STAT4 occurs mainly in lymphoid and myeloid cells, but also vascular endothelial cells (Torpey et al., 2004), thus, for 018 to modulate this pathway, VACV would need to infect these cell types *in vivo*. As the 018-binding interface is conserved between STAT1 and 4, functionally, 018 could prevent STAT4 recruitment to active receptors. Whether the 018:STAT4 interaction plays a physiological role during infection remains to be determined.

As STATs are an important class of drug targets in a wide range of diseases (Miklossy et al., 2013), the highly specific interaction of 018 for STAT1 and 4, notably via the HxH motif, presents a potential avenue for development of STAT1/4-selective inhibitors via rational drug design. One approach could involve the construction of peptidomimetic derivatives of the HxH motif in a structure-guided manner to deliver high-affinity binders.

The STAT1-binding region of 018 possesses significant sequence similarity to the STAT1- binding region of V/W and P proteins from NiV, a paramyxovirus first discovered in Malaysia in 1998 (Chua, 2000). NiV is highly pathogenic and has caused numerous sporadic outbreaks, 19 including recently in Kerala, India (Arunkumar et al., 2019) and no effective treatments or vaccines are available (Hauser et al., 2021). Previous studies of NiV-V showed it sequesters STAT1 and 2 and prevents STAT1 phosphorylation (Rodriguez et al., 2002). Our data advance this observation and show that in the context of IFN-II signalling, the NiV STAT1 binding region can block STAT1:pIFNGR1 association. Mechanistically, this is most relevant to the V and P proteins due to their cytoplasmic location (Shaw et al., 2004). Although W harbours an identical STAT1-binding region, it traffics STAT1 to the nucleus to prevent STAT1 activation (Shaw et al., 2004). As we anticipate that the 114-VVYHDHGG-121 region of NiV-V/W and P bind in an analogous fashion to the 14-VFIHGHDG-19 of 018, the 018:STAT1 structure can aid understanding of previous mutagenesis studies of the NiV STAT1 binding region (Ciancanelli et al., 2009; Hagmaier et al., 2006; Jensen et al., 2020; Ludlow et al., 2008; Satterfield et al., 2019).

Intrinsically disordered proteins that harbour short linear motifs (SLiMs), such as the STAT1- binding region from NiV and 018, are important mediators of virus-host interactions (Mishra et al., 2020). SLiMs are advantageous to viruses because they offer high flexibility and typically can evolve at fast rate allowing quick adaptation to changing host environments (Xue et al., 2014). Virus SLiMs that mimic eukaryotic linear motifs have appeared as a prevalent virus strategy to hijack cellular machinery and disable host defences (Davey et al., 2011; Hagai et al., 2014; Lasso et al., 2021). Because SLiMs are short and evolve easily, they have emerged predominantly independent of their host mimics rather than by horizontal gene transfer from the host (Elde and Malik, 2009; Hagai et al., 2014). In the context of the 018/NiV-V STAT1- binding motif, although it is possible cellular proteins do exist that bind STAT SH2 domains in pTyr-independent manner, none have been identified and thus 018/NiV-V might not mimic a cellular interaction. The STAT1-binding motif described here likely represents a striking example of convergent evolution in diverse virus families and has produced an unconventional binding mechanism to target STAT1. Consistent with the notion that SLiMs preferentially target proteins central to multiple networks (Dyer et al., 2008), STAT1 is required for ISG induction in response to all IFN families (IFN-I, II and III). The existence of the shared motif between disparate viruses highlights its importance as an efficient moiety for inhibiting IFN-induced signalling.

It is notable that amongst the different STAT domains, the SH2 domain is the most highly conserved across various species (Park et al., 2008). Targeting this conserved domain may enable these viral antagonists to function in multiple species and contribute to the broad species specificity of VACV and NiV. Consistent with this notion, data presented here show that 018 antagonises human IFN signalling and contributes to VACV virulence in mice. Along the same lines, the VACV decoy IFN receptors B8 and B18 bind and neutralise IFNs from many species (Alcamí and Smith, 1995; Symons et al., 1995).

Poxviruses encode multiple antagonists of IFN-induced signalling. The earliest functioning of these, the viral phosphatase vH1, is carried within virions and released into the cytoplasm upon infection where it might dephosphorylate STAT1, although this activity has been demonstrated only in vitro (Najarro et al., 2001; Schmidt et al., 2013). Multiple reports have shown that shortly after VACV infection cells are refractory to pSTAT1 activation by IFN-II stimulation (Mann et al., 2008; Najarro et al., 2001; Schmidt et al., 2013). Hitherto, this phenotype was mainly attributed to vH1, however deletion of 018 led to an almost complete rescue of pSTAT1 levels despite the presence of vH1, demonstrating that during infection 018, rather than vH1, is responsible for this phenotype. Consistent with this early block, 018 is one of the earliest viral proteins detected during infection (Soday et al., 2019).

Despite apparent redundancy in inhibition of IFN-induced signalling by VACV, deletion of individual IFN antagonists leads to virus attenuation *in vivo* (Figure 4; Symons et al., 1995; Unterholzner et al., 2011). These non-redundant phenotypes may stem from each inhibitor having different locations or expression kinetics or being multifunctional. Unlike intracellular inhibitors, B18 and B8 are secreted from cells and thus can neutralise IFNs extracellularly and distally. Also, B18 can bind to cell surface glycosaminoglycans and thereby inhibit IFN-I- induced signalling in uninfected cells (Alcamí et al., 2000; Montanuy et al., 2011). This is the major mechanism by which B18 contributes to virulence (Hernáez et al., 2018). Although the B18 orthologue from Yaba-like disease virus (a yatapoxvirus) binds and inhibits IFN-III, the VACV B18 protein does not (Huang et al., 2007), and there is no known specific inhibitor of IFN-III-induced signalling made by VACV. Nonetheless, cells infected with VACV are refractory to pSTAT1 induction after IFN-III stimulation, and IFN-III expression during viral infection has little effect on VACV replication (Bandi et al., 2010; Bartlett et al., 2005). These observations may be explained by the action of 018. Differences in expression kinetics of VACV IFN antagonists could also affect redundancy, for, although B18 functions upstream of 018, VACV lacking 018 showed enhanced levels of pSTAT1 after IFN-I stimulation. Lastly, virus proteins are often multifunctional. Indeed VACV protein C6, which inhibits both IFN production (Unterholzner et al., 2011) and IFN-I signalling in the nucleus (Stuart et al., 2016) also degrades HDAC4 and 5, which are restriction factors for VACV, and this might be the major factor contributing to virulence (Lu et al., 2019; Soday et al., 2019).

Deletion of IFN antagonists can improve the safety and immunological memory of VACV- based vaccine vectors (Albarnaz et al., 2018). Modified vaccinia Ankara (MVA) is a widely used VACV-based vaccine vector and expresses 018 (Wennier et al., 2013). MVAs expressing SARS-CoV-2 proteins have been described as potential vaccine candidates, and thus our findings can inform further development (Chiuppesi et al., 2020; García-Arriaza et al., 2021; Liu et al., 2021).

In summary, we describe a viral mechanism to antagonise IFN-induced signalling by occupancy of the STAT1 SH2 domain to prevent STAT1 receptor association and subsequent ISG expression. The structure of VACV protein 018 complexed with STAT1 illustrates how a viral protein has evolved an unconventional strategy to bind an SH2 domain with high affinity. The biological importance of 018 is shown by its contribution to virus virulence despite the presence of other IFN antagonists. Finally, this study highlights how disparate viruses can evolve highly similar motifs to target a host response that poses a common threat to all viruses.

## Acknowledgments

This work was supported by a Wellcome Trust Principal Research Fellowship (090315, to G.L.S.). C.T.C was funded by the BBSRC Doctoral Training Partnership. T.P. was funded by the MRC Doctoral Training Partnership. M.T.A. was funded by a Harry Smith vacation studentship from the Microbiology Society. J.P.S., C.R.C. and H.D.H. were supported by the NIAID Division of Intramural Research. We thank Diamond Light Source for access to macromolecular crystallography beam line i04 (proposal 25402). We are grateful for access to instrumentation and support by the X-ray crystallographic and Biophysical research facilities at the Department of Biochemistry, University of Cambridge.

Henrietta Lacks, and the HeLa cell line that was established from her tumour cells without her knowledge or consent in 1951, have made significant contributions to scientific progress and advances in human health. We are grateful to Henrietta Lacks, now deceased, and to her surviving family members for their contributions to biomedical research.

## Author contributions

Conceptualisation: C.T.C., T.P. and G.L.S.

Methodology: C.T.C., T.P. and J.P.S.

Formal Analysis: C.T.C., T.P. and J.P.S.

Investigation: C.T.C., T.P., J.P.S., C.R.C. and M.T.A.

Resources: T.P., M.H., H.D.H. and G.L.S.

Data Curation: C.T.C, T.P., J.P.S. and M.H.

Writing: Original draft: C.T.C. and T.P.

Writing: Reviewing and Editing: C.T.C, T.P., J.P.S., C.R.C., M.T.A., M.H., H.D.H., and G.L.S.

Visualisation: C.T.C. and T.P.

Supervision: M.H., H.D.H. and G.L.S.

Project Administration: C.T.C. and G.L.S.

Funding Acquisition: M.H., H.D.H. and G.L.S.

## STAR methods

### KEY RESOURCES TABLE

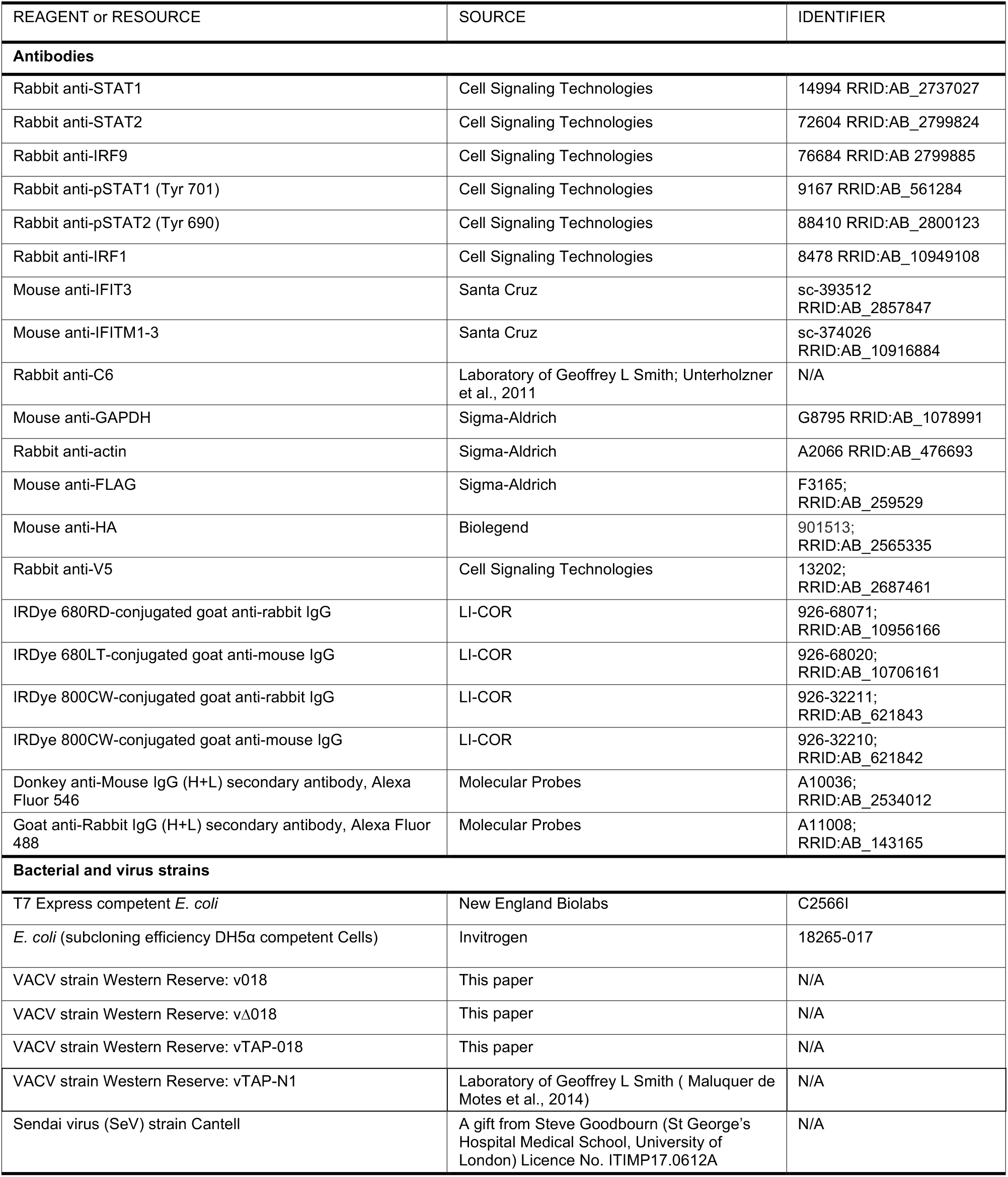
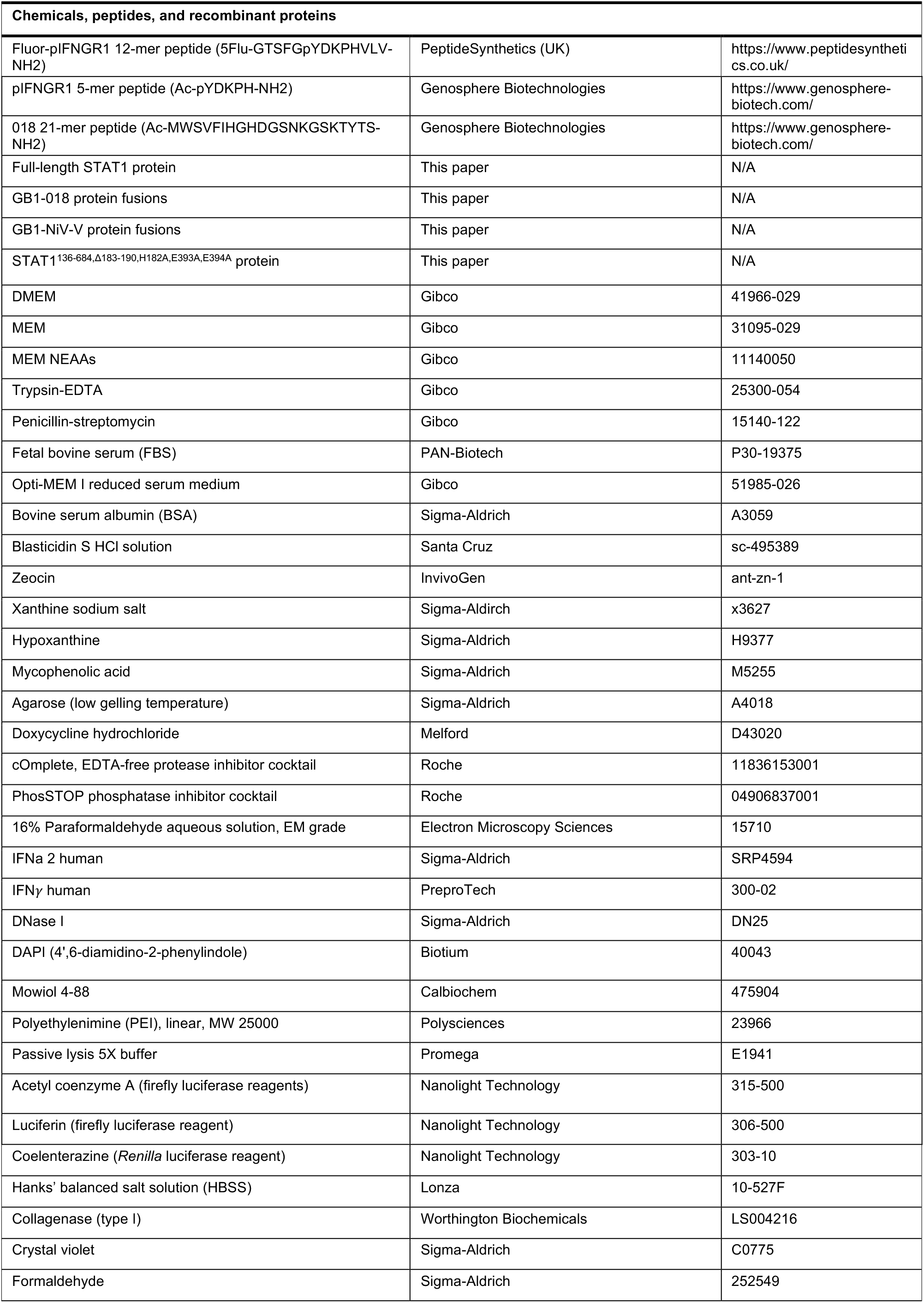
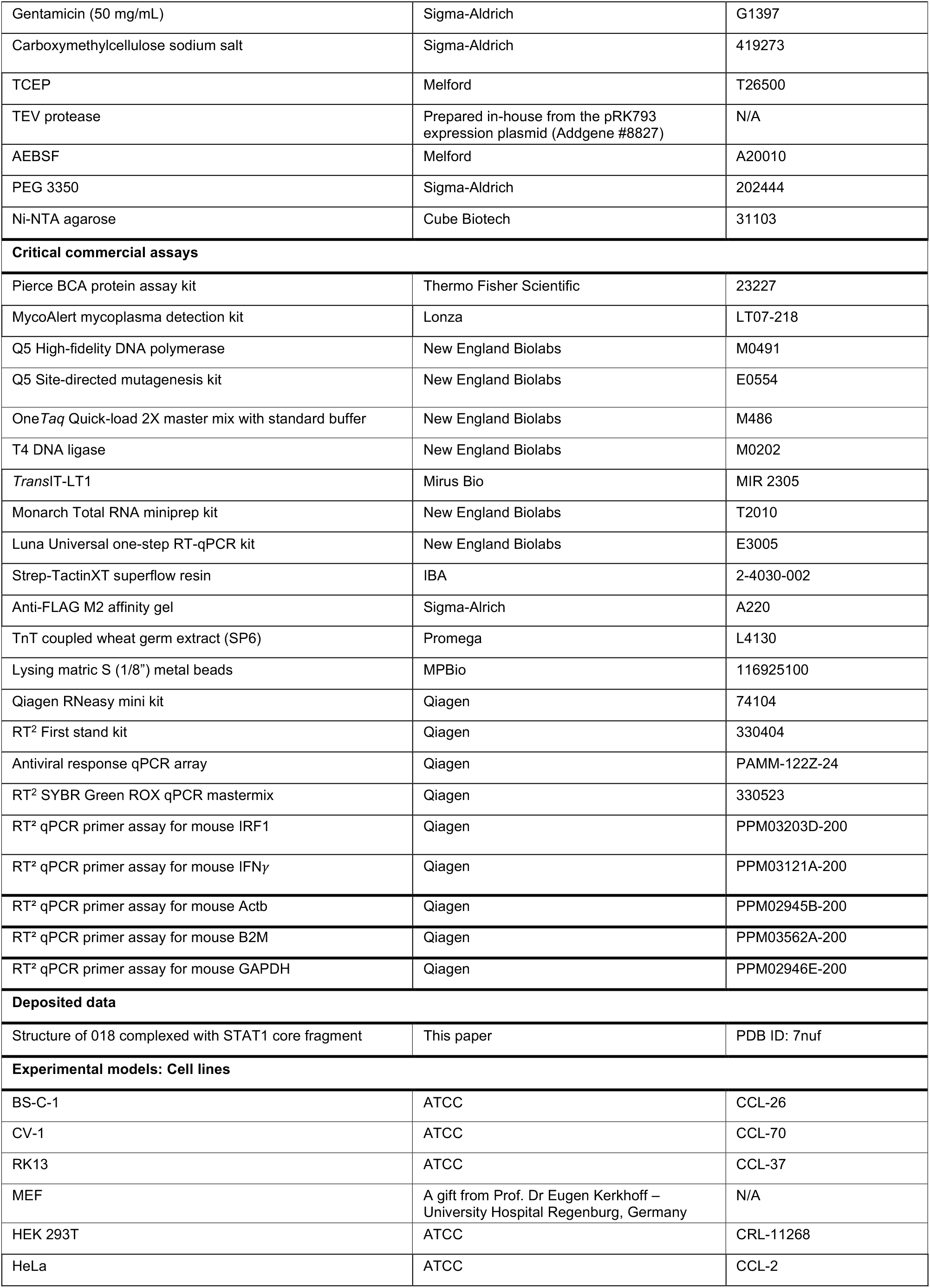
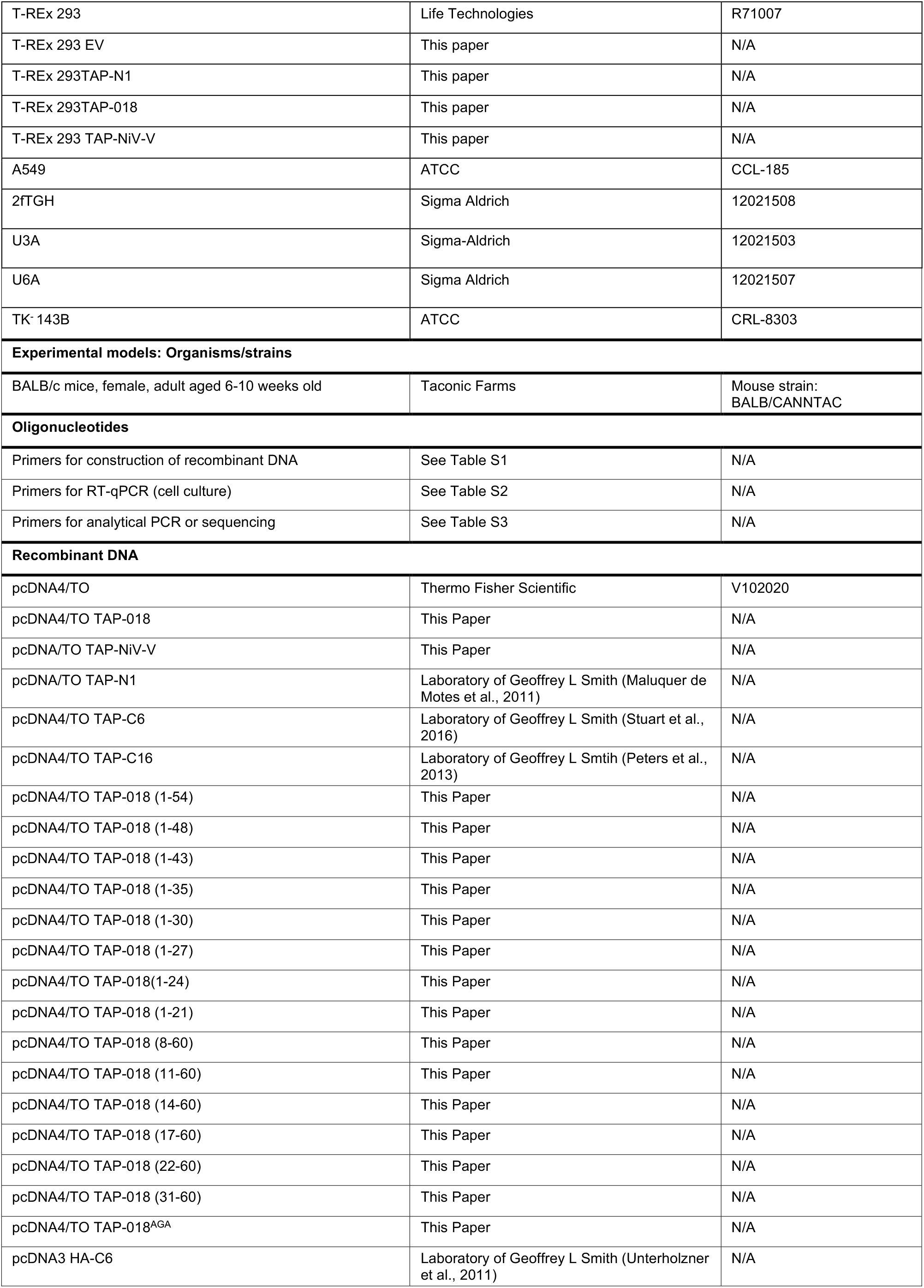
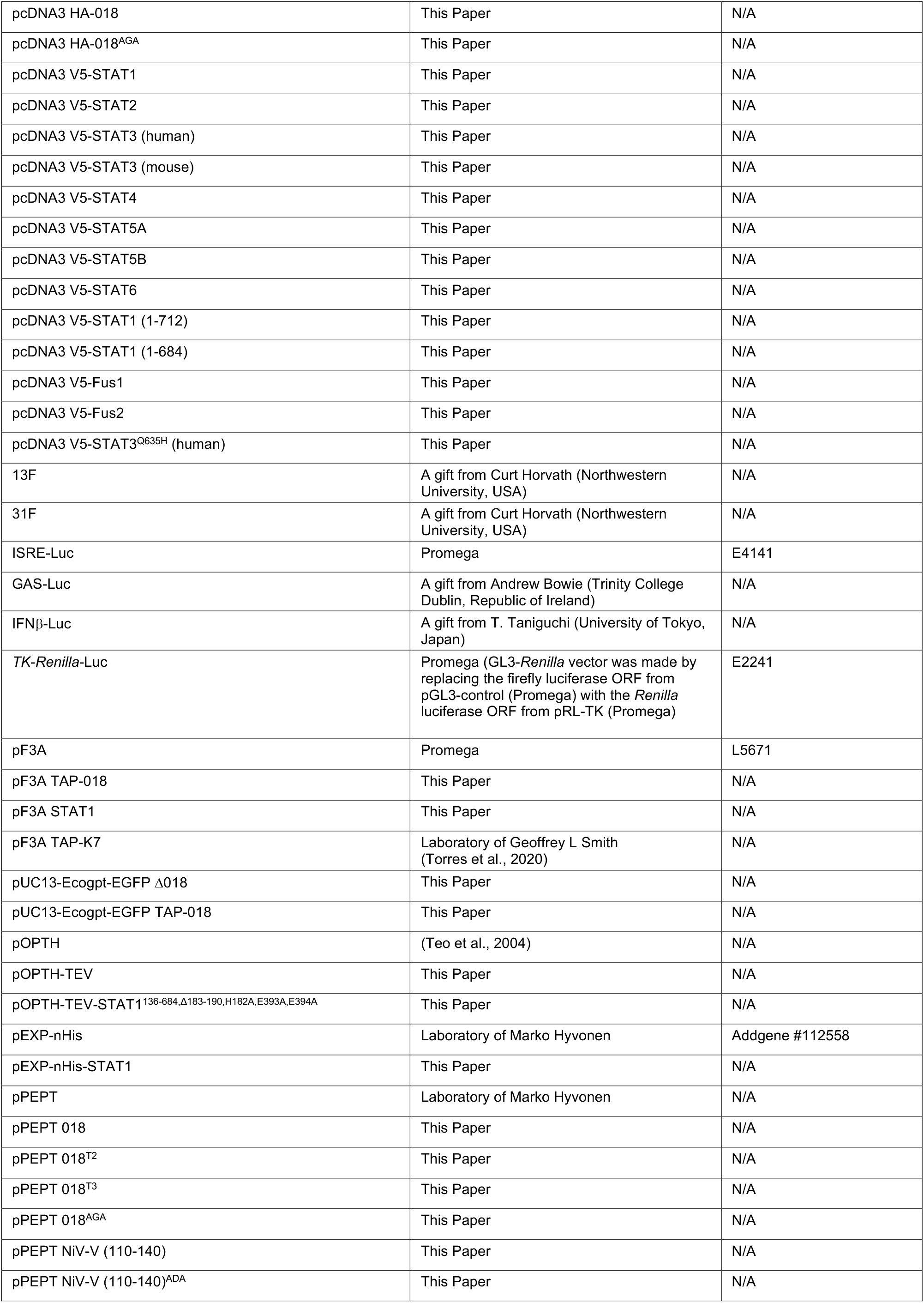
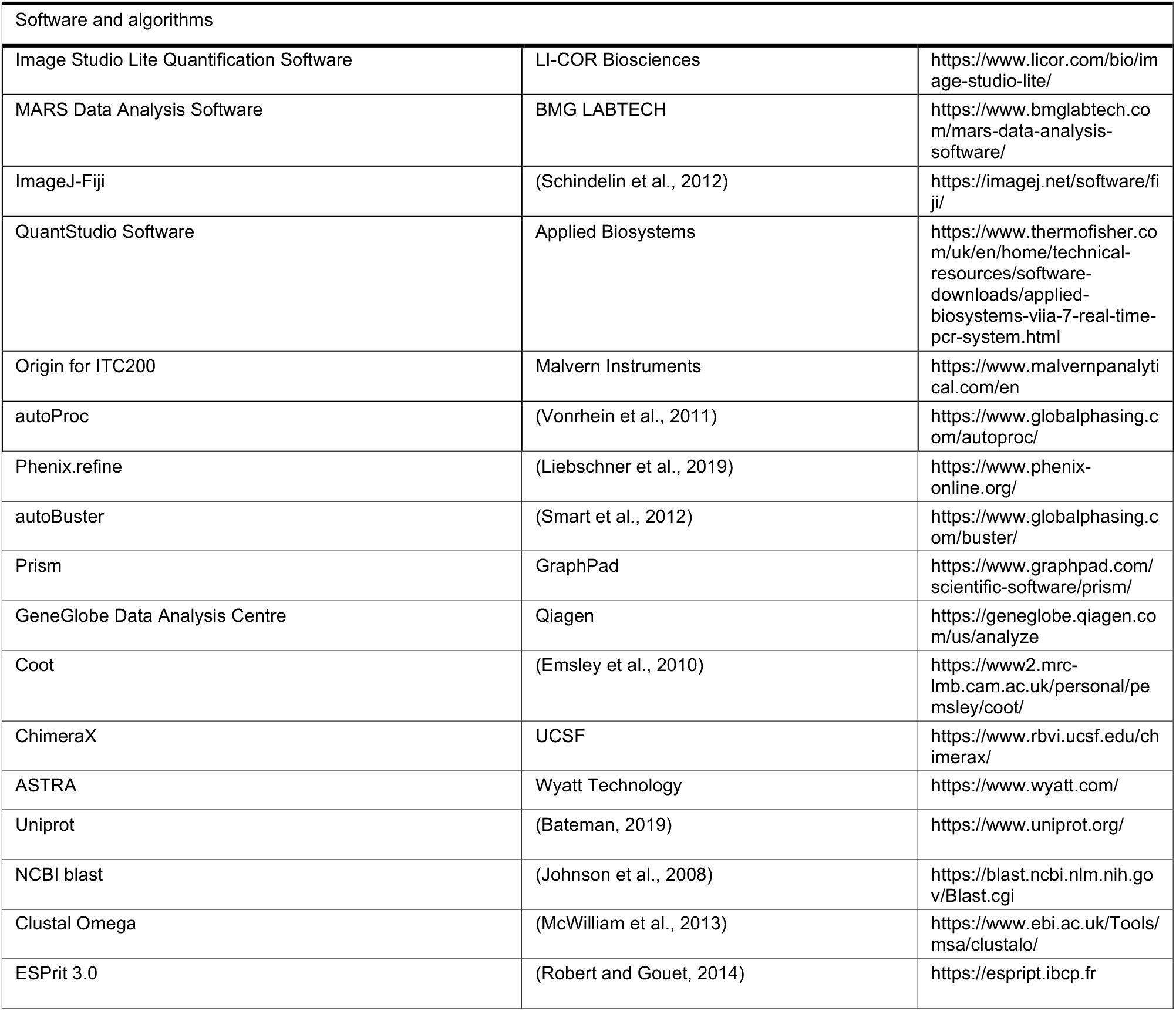

### RESOURCE AVAILABILITY

#### Lead contact

Further information and requests for resources and reagents should be directed to and will be fulfilled by the lead contact, Geoffrey L Smith.

#### Materials availability

See above.

#### Data and code availability

018:STAT1 X-ray crystallographic structure has been deposited on the PDB under the accession code 7nuf.

### EXPERIMENTAL MODEL AND SUBJECT DETAILS

#### Cell lines

BS-C-1 (ATCC), CV-1 (ATCC), MEFs (A gift from Prof. Dr Eugen Kerkhoff), HEK 293T (ATCC), A549 (ATCC), 2fTGH (a human cell line containing the selectable marker guanine phosphoribosyltransferase regulated by IFN-α, Sigma-Aldrich), U3A (a 2fTGH derived STAT1^-/-^ cell line, Sigma-Aldrich), U6A (a 2fTGH derived, STAT2^-/-^ cell line, Sigma-Aldrich), and TK^-^ 143B cells (ATCC) were maintained in DMEM (Gibco) supplemented with 10% fetal bovine serum (FBS, PAN-Biotech) and penicillin-streptomycin (PS, 50 μg/mL, Gibco). T-REx 293 cells (Life technologies) were maintained in DMEM supplemented with 10% FBS, PS (50 μg/mL) and blasticidin (10 μg/mL, Santa Cruz), and T-REx 293 derived cells lines expressing EV, TAP-N1, TAP-018, or TAP-NiV-V were further supplemented with zeocin (100 ug/mL, Invivogen). HeLa (ATCC) and RK13 cells (ATCC) were maintained in MEM (Gibco) supplemented with 10% FBS, PS (50 μg/mL) and 1% non-essential amino acids (Gibco). The construction of T-REx 293 cell lines expressing proteins inducibly is outlined in the methods detail section.

#### Viruses

Recombinant vTAP-N1 was derived from VACV strain WR (VACV-WR, GenBank: AY243312.1) (Maluquer de Motes et al., 2014). v018, vΔ018 and vTAP-018 described in this paper were all derived from VACV strain WR and their construction is outlined in the methods detail section.

#### Animals

Specific pathogen-free BALB/c mice were obtained from Taconic Farms. 6-10 weeks old female mice were used in all experiments. Mice were housed under specific pathogen-free conditions (including negativity for murine norovirus, mouse parvovirus, and mouse hepatitis virus) and were maintained on standard rodent chow and water supplied *ad libitum*. All animal studies were approved by and performed in accordance with the Animal Care and Use Committee of the National Institute of Allergy and Infectious Diseases, USA.

### QUANTIFICATION AND STATISTICAL ANALYSIS

Significances were calculated in Prism (GraphPad) by either Dunnett’s T3 multiple comparisons test or Unpaired t-test with Welch’s correction as indicated. For anti-viral array data (**Figure 4G**), analysis and Unpaired t-tests was performed using GeneGlobe Data Analysis Centre (Qiagen). All significances are indicated with P values.

### METHOD DETAILS

#### Orthologue alignments

Identifiers for poxvirus genomes from which the amino acids sequences of 018 orthologues were derived: vaccinia strain Western Reserve (VACV-WR, GenBank: AY243312.1), Modified vaccinia Ankara (MVA, GenBank: AY603355.1), variola virus (VARV, GenBank: X69198.1), monkeypox virus (MPXV, GenBank: AF380138.1), cowpox virus strain Brighton Red (CPXV-BR, GenBank: AF482758.2), ectromelia virus (ECTV, GenBank: AF012825.2), camelpox virus (CMLV, GenBank: AF438165.1), rabbitpox virus (RPXV, GenBank: AY484669.1), raccoonpox virus (RCNV, GenBank: KP143769.1), skunkpox virus (SKPTV, GenBank: KU749310.1), taterapox virus (TATV, GenBank: DQ437594.1), Cotia virus (COTV, GenBank: HQ647181.2), Yoka poxvirus (YKPV, GenBank: HQ849551.1). Alignments were performed using Clustal Omega (McWilliam et al., 2013) and conservation annotation was performed using ESPirt 3.0 (Robert and Gouet, 2014).

#### Plasmids

The 018 open reading frame was codon-optimised for expression in human cells and was synthesised by Gene Art (Thermo Fisher Scientific). All plasmids are described in recombinant DNA key resource table and primers used for construction in **Table S1**.

#### Construction of T-REx 293 cell lines expressing proteins inducibly

T-REx 293 were transfected with pcDNA4/TO expression plasmids (pcDNA4/TO (EV), pcDNA4/TO TAP-018, pcDNA4/TO TAP-N1 and pcDNA4/TO TAP-NiV-V) using Transit- LT1 (Mirus Bio). Prior to transfection, pcDNA4/TO expression plasmids were linearised using *PvuI* (NEB) to decrease the potential for plasmid-chromosomal integration within the viral ORF. Cells with integrated plasmid were selected in the presence of blasticidin (10 μg/mL) and zeocin (100 μg/mL) and single clones were obtained by limiting dilution. Clones were amplified and lysates were analysed for the expression of TAP-tagged protein by immunoblotting.

The expression of TAP-tagged proteins from T-REx 293-derived cells was induced by addition of doxycycline (100 ng/mL, Santa Cruz) to the medium for 24 h for all experiments.

#### Construction of recombinant VACVs

Recombinant VACVs (vΔ018 and vTAP-018) were constructed using transient dominant selection (Falkner and Moss, 1990). To construct the pUC13_Ecogpt_EGFP_Δ018 plasmid to remove the entire 018 ORF, the downstream (301 bp) and upstream (300 bp) flanking regions of the 018 ORF were amplified by PCR from purified VACV (strain WR) DNA. A 15 bp complementary sequence was added to the internal upstream and downstream primers to facilitate joining of the two-flanking regions by overlapping PCR. The resulting PCR product was then ligated into pUC13- Ecogpt-EGFP using *Pst*I (NEB) and *Bam*HI (NEB) cloning sites. To construct the pUC13-Ecogpt-EGFP TAP-018 plasmid, the downstream flanking and 018 ORF (484 bp) and the upstream region of the ORF (300 bp) were amplified by PCR separately from purified VACV (strain WR) DNA. The 018 ORF plus downstream flanking region PCR product was ligated into pcDNA4/TO vector containing an N-terminal TAP-tag using *Not*I (NEB) and *Xba*I (NEB) as an intermediate cloning step. The N-terminal TAP tag fused 018 ORF + downstream flanking region was then amplified by PCR using primers that added a 20- bp overhang sequence complementary to the upstream flanking PCR product. The two PCR products were then joined by overlapping PCR and the product was ligated into pUC13_Ecogpt_EGFP using *Pst*I and *Bam*HI cloning sites.

To construct vΔ018, CV-1 cells were seeded in a T-25 flask to be 70 % confluent the following day. CV-1 cells were then infected with VACV (strain WR) at 0.05 pfu/cell and after 1 h 30 min, the inoculum was removed and cells were transfected with 7.5 μg of pUC13_Ecogpt_EGFP_Δ018 using Transit-LT1 (Mirus Bio). Two days p.i., the majority of cells displayed cytopathic effect and were harvested by scrapping cells into the culture medium. Cells were sedimented by centrifugation (500 RCF) and resuspended in 0.5 mL of infection medium (DMEM supplemented with 2% FBS and PS (50 μg/mL). Samples were freeze-thawed three times to lyse cells and release progeny virus and sonicated to disperse particulate material. A series of progeny virus dilutions (10^-1^, 10^-2^ and 10^-3^ diluted in infection medium) were used to inoculate BS-C-1 cells in 6-well plates that had been preincubated in infection medium, supplemented with mycophenolic acid (25 μg/mL; MPA, Sigma-Alrich), xanthine (250 μg/mL; X, Sigma-Alrich) and hypoxanthine (15 ug/mL; HX, Sigma-Alrich) for 24 h. After 1 h 30 min, the inoculum was removed and replaced with a MEM, 1% (w/v) low gelling temperature agarose (Sigma Alrich), supplemented with MPA, HX, and X. After three days, EGFP-expressing plaques were picked, representing virus that had integrated the pUC13- Ecogpt-EGFP-Δ018, and then further plaque purified in the absence of MPA, HX and X. The genotype of these plaques was then determined by PCR using primers that flank the 018 ORF (**Table S3**) and VACVs containing the desired mutation (vΔ018) or wild type genotype were isolated. vTAP-018 was produced using the same strategy as described above, except that vΔ018 was used as the parental VACV into which the TAP-018 ORF was inserted at its natural locus. Stocks of VACVs were grown in RK13 cells and titrated by plaque assay on BS-C-1 cells.

#### Purification of VACVs by sedimentation through sucrose

VACVs were purified by two rounds of ultracentrifugation through a sucrose cushion as described (Joklik, 1962) and stocks were resuspended in 1 mM Tris-HCl pH 9.0 for cell culture work or in Hank’s balanced salt solution (HBSS) + 0.1% BSA for *in vivo* work. Virus titres were determined by plaque assay.

Viral DNA for vΔ018 and wild-type sibling virus v018 was isolated from sucrose purified virus stocks by phenol:chloroform extraction. Whole genome sequencing of viruses was performed by MircobesNG and virus sequences were aligned to VACV strain WR reference genome (VACV-WR, GenBank: AY243312.1).

#### VACV infection for cell culture

Virus inoculums were prepared in DMEM supplemented with 2% FBS (infection medium) and virus adsorption was performed at 4 ^⁰^C for 1 h with gentle agitation every 10 mins. At time 0 h p.i., virus inoculum was removed and replenished with infection medium, and infection was continued at 37 ^⁰^C.

#### Virus growth and spread assays

For virus growth curves, BS-C-1 cells were grown to confluence in T-25 flask then infected at 5 pfu/cell. At 1, 8 and 24 h p.i., extracellular and cell-associated virus were harvested by collecting either the supernatant or cell monolayers, respectively. Supernatants were cleared by centrifugation (21,000 RCF) to remove detached cells and debris. Cell monolayers were scrapped into new medium and subjected to three cycles of freeze-thawing followed by sonication to release intracellular virus. Viral titres were determined by plaque assay.

Virus spread was determined by analysis of plaque size growth. Confluent BS-C-1 and RK13 cells in 6-well plates were infected with 30 pfu/per well. At 1 h p.i., medium was replaced with a semi-solid MEM overlay supplemented with, L-glutamine, 2% FBS and 1.5 % (w/v) carboxymethylcellulose (Sigma-Aldrich). At 72 h p.i., the semi-solid overlay was removed, and monolayers were stained with crystal violet (Sigma-Aldrich).

#### Immunoblotting

For immunoblotting analysis, cells were washed once in chilled PBS and harvested on ice by scrapping into lysis buffer (Tris pH 8.0, 150 mM NaCl and 1% NP-40, supplemented with protease (cOmplete Mini, Roche) and phosphatase inhibitors (PhosSTOP, Roche). Cell lysates were incubated with rotation at 4 ^⁰^C for 15 mins before being cleared by centrifugation (21,000 RCF) and protein concentrations were determined using BCA Protein Assay (Pierce). Lysates were mixed with 4X SDS-gel loading buffer and incubated at 100 ^⁰^C for 5 min to denature protein. Samples were briefly centrifuged (17,000 RCF) before loading onto either SDS-polyacrylamide gels or NuPAGE (4 to 12%, 1 mm, Bis-Tris gels (Thermo Fisher Scientific) along with protein ladder (Abcam) and separated by electrophoresis. Protein gels were incubated in transfers buffer (25 mM Tris, 250 mM glycine, 20% (v/v) methanol) with agitation for 15 min. Gels were transferred onto a nitrocellulose transfer membrane (0.2 μM pore size, GE Healthcare) using a semi-dry transfer system (Trans-tubro blot, BioRad). Nitrocellulose membranes were allowed to dry for 30 mins and then blocked with 5% (w/v) BSA (Sigma), in TBS containing 0.1% Tween-20 for 1 h at room temperature (RT). Primary antibodies (see Key Resources Table) were diluted in blocking buffer and incubated with membranes overnight at 4 ^⁰^C. Membranes were probed with fluorophore-conjugated secondary antibodies (LI-COR Biosciences) diluted in 5% (w/v) non-fat milk in PBS containing 0.1 % (v/v) Tween-20 and incubated at RT for 45 min. Membranes were imaged using the Odyssey CLx imagining system (LI-COR Biosciences). Protein band intensities were quantified using Image Studio software (LI-COR Biosciences).

#### Reporter gene assays

HeLa cells (for GAS-Luc reporter) or HEK 293T cells (for ISRE-Luc and IFNβ-Luc) in 96- well plates were co-transfected with 75 ng of firefly luciferase reporter (GAS-Luc, ISRE-Luc or IFNβ-Luc), 10 ng of *TK-Renilla* plasmid and the desired expression plasmid using Trans- LT1 (Mirus Bio). In cases where different doses of the expression plasmids were used, the lower doses were topped up by addition of EV plasmids so that equal amounts of DNA were transfected per well. Twenty-four h post transfection, cells were either non-stimulated, or stimulated with IFNα (1000 U/mL, Sigma-Aldrich), IFNγ (25 ng/mL, PreproTech) or SeV (a gift from Steve Goodbourn) for 6, 8 or 24 h, respectively. Following stimulation, cells were lysed in passive lysis buffer (Promega) and firefly and *Renilla* luciferase luminescence were measured using a FLUOstar luminometer (BMG). Firefly values were normalised to *Renilla* luciferase readings and fold inductions were calculated for each sample relative to their own non-stimulated values. Results are presented as individual data point without P values. Relative protein expression levels were determined by immunoblotting.

#### Immunofluorescence

For VACV infection, A549 cells were seeded onto sterile glass coverslips (Thickness no. 0.13- 0.17 mm, diameter 19 mm, Thermo Fisher Scientific) in 12-well plates. Twenty-four h after seeding, cells were serum starved for 16 h prior to infection. Cell were infected at 10 pfu/cell and at 2 h p.i., cells were washed once in medium before being stimulated with IFNγ (25 ng/mL, PreproTech) for 30 min.

For transfection, HeLa cells were seeded onto sterile glass coverslips (Thickness no. 1.5, diameter 22 mm, Thermo Fisher Scientific) in 6 well plates. Twenty-four h after seeding, cells were transfected with 0.8 μg of expression plasmids using TransIT LT1 (Mirus Bio). Five h post transfection, medium on cells was replaced with serum-free medium to serum starve cells for 16 h. Cells were then either non-stimulated, stimulated with IFNα (1000 U/mL, Sigma- Alrich) or stimulated with IFNγ (25 ng/mL, Prepotech) for 1 h. Following stimulation, cells were fixed in 8% (v/v) paraformaldehyde (PFA, Electron Microscopy Sciences) in 250 mM HEPES pH 7.5 for 5 min on ice followed by 25 min at RT. After fixation, cells were incubated for 5 min with 50 mM ammonium chloride in PBS to quench free aldehydes. Cells were permeabilised by incubating with ice-cold, 100 % methanol at -20 ^⁰^C for 10 min. Cells were blocked in IF buffer (10 % v/v FBS in PBS) for 30 min before staining with primary antibodies for 1 h. After washing, coverslips were then stained with secondary antibodies (AlexPhore) diluted in IF buffer, supplemented with 5% (v/v) serum from primary antibody source animal for each secondary antibody for 30 mins. Coverslips were then mounted using Mowiol 4-88 containing 4’,6-diamidino-2-phenylindole (DAPI) on to microscope slides (Menzel-Gläser). Slides were visualised with a LSM 780 inverted confocal microscope (Zesis) and images were processed using Image J (Schindelin et al., 2012).

#### RT-qPCR

A549 cells in 12-well plates were infected at 10 pfu/cell. At 2 h p.i. cells were washed once in medium before being stimulated with IFNγ (25 ng/mL, PreproTech) for 1 h. Following stimulation, RNA was harvested using Monarch Total RNA Miniprep Kit (NEB) according to manufacturer’s instructions including an optional on-column genomic DNA digestion step. RT-qPCR was performed using Luna Universal One-Step RT-qPCR Kit (NEB). Oligonucleotide primers (**Table S2**) targeting HRPT and IRF1 were designed using PrimerQuest Tool (IDT). RT-qPCRs were carried out using a real-time PCR system (Thermo Fisher Scientific) and fold-inductions of ISG levels were calculated using 2^−ΔΔCt^ taking mock non-stimulated readings as the basal level sample and HRPT as the control housekeeping gene.

#### Pulldowns

For infection, BS-C-1 cells or MEFs in T-25 flasks were infected at 10 pfu/cell with either vTAP-018 or vTAP-N1. For transfection, 2fTGH, U3A, U6A and HEK 293T cells in 10-cm dishes, were transfected using either TranIT LT1 (Mirus Bio) for 2fTGH, U6A and U3A cells or polyethylenimine (PEI, 2 μl of 1 mg/mL stock per μg of DNA, Polysciences) for HEK 293T cells. Prior to transfection, medium was replaced with DMEM supplemented with 2% FBS. At 12 h p.i. or 18 h after transfection, cells were lysed in Tris-based IP buffer (50 mM Tris pH 7.4, 150 mM NaCl, 0.5% (v/v) NP-40) supplemented with protease (cOmplete Mini, Roche) and phosphatase inhibitors (PhosSTOP, Roche). Cell lysates were incubated with rotation at 4 ^⁰^C for 15 mins before being cleared by centrifugation (21,000 RCF). A fraction of cleared lysate was taken for input samples and the remaining lysate was incubated with 30 μl of one of the following affinity resins washed and equilibrated in IP buffer: (i) Strep-Tactin XP super flow (IBA) for pulldown of TAP-tagged proteins via Strep-tag II epitope; (ii) anti-FLAG M2 affinity gel (Sigma-Aldrich) for immunoprecipitation of either FLAG or TAP-tagged protein via the FLAG epitope. Samples were incubated with affinity resins at 4 ^⁰^C with rotation for 1 h 30 min. Samples were washed three times in IP buffer and proteins were eluted from beads by addition of 2X SDS-gel loading buffer. Subsequently, samples were analysed by either Nu-PAGE (Thermo Fisher Scientific) or SDS-PAGE followed by immunoblotting.

For pulldowns using proteins produced from a cell-free transcription and translation system, the TnT Sp6 High-Yield wheat germ protein expression system (Promega) was utilised according to manufacturer’s instructions.

#### *In vivo* experiments

Female BALB/c mice 6-10 weeks old were anesthetized and infected intranasally (i.n.) with 10^3^ pfu for measurement of weight change and pulmonary virus titres or 10^5^ pfu for extraction of RNA and RT-qPCR experiments. A final inoculation volume of 20 μl (10 μl per nostril) was used with VACVs diluted in HBSS + 0.1% BSA to achieve the required dose. The actual dose administered was confirmed by plaque assay of the diluted virus inoculum.

For weight change experiments, mice were weighed daily. For virus titration experiments, lungs were collected at 3, 7 and 9 days p.i. and single-cell suspensions were prepared by chopping with scissors followed by collagenase I digestion (Worthington Biochemicals) for 60 mins at 37 ⁰C. Cells were disrupted by vigorous pipetting and suspensions were freeze-thawed three times to release virus and infectious virus titres were determined by plaque assay on TK^-^ 143B cells. For RT-qPCR experiments the upper lobes of lungs were removed and immediately placed in buffer RLT (Qiagen). Lungs were homogenized and RNA was isolated using Lysing Matrix S (1/8^”^) metal beads (MPBio) and a FastPrep^®^-24 Instrument (MPBio). RNA was then purified using a Qiagen RNeasy Mini Kit (Qiagen). An on-column DNAse (Sigma-Aldrich) digestion was performed prior to RNA elution. cDNA was synthesised using the RT^2^ First Strand Kit (Qiagen) with ∼ 1.2 g of RNA/sample. cDNA was then loaded onto an Antiviral Response qPCR array (Qiagen) or onto a separate plate for the analysis of IRF1 and IFNγ for which individual RT^2^ qPCR primer assays (Qiagen) were obtained. RT-qPCRs were carried out using RT^2^ SYBR Green ROX qPCR mastermix (Qiagen) and a real-time PCR system (Thermo Fisher Scientific) and fold-changes of genes were calculated by comparing Ct values of individual vΔ018-infected mice (n=4) to the Ct averages of v018-infected mice (n=3) using the 2^−ΔΔCt^ method. Fold changes of genes were normalised against 5 standard housekeeping genes included on the Antiviral Response qPCR array (Qiagen) or against 3 standard housekeeping genes (Actb, B2M and GAPDH, Qiagen) for analysis of IRF1 and IFNγ (Qiagen). Data analysis and significances were performed using manufacturer’s software (GeneGlobe Data Analysis Centre, Qiagen).

#### Protein expression and purification

The purity of protein preparations was analysed by SDS-PAGE and subsequent Coomassie blue staining (**Figure S7**).

Full-length STAT1 and STAT1^136-684,Δ183-190,H182A,E393A,E394A^ expression plasmids were transformed into *E. coli* T7 Express cells (NEB) and plated overnight on LB agar supplemented with 100 μg/mL of ampicillin. The next day colonies of transformed cells were collected and used to inoculate 1 L (TB) medium supplemented with 100 μg/mL of ampicillin and were grown at 37 ⁰C in 2 L flasks until OD600 of 0.8-1.2. Cultures were cooled to 18 ⁰C and incubated overnight with 0.4 mM IPTG to induce protein expression. Cells were collected by centrifugation and resuspended in lysis buffer (25 mL of 50 mM Tris-HCl, pH 8.0, 300 mM NaCl, 20 mM imidazole, 1 mM AEBSF, 1 mM TCEP) and lysed by sonication. Cell lysates were centrifugated at 40,000 RCF for 30 min and the cleared supernatant was loaded on a 3 mL Ni-NTA agarose resin (Cube Biotech) or on a 5mL HisTrap HP column (Cytiva). The column matrix was washed with 10 column volumes (CV) of wash buffer (50 mM Tris-HCl, pH 8.0, 300 mM NaCl, 20 mM imidazole, 1 mM TCEP). Proteins were eluted with 50 mM Tris-HCl, pH 8.0, 300 mM NaCl, 200 mM imidazole, 1 mM TCEP into 2 mL fractions. Fractions containing the proteins of interest were pooled. STAT1^136-684, Δ183-190, H182A, E393A, E394A^ fractions were incubated with 100 μL of 2 mg/mL TEV protease (prepared in-house) overnight at 4 ⁰C to remove the N-terminal His6 affinity tag. STAT1 proteins were then diluted ten-fold in heparin buffer A (20 mM Tris-HCl, pH 8.0, 1 mM EDTA) and loaded on a 5 mL HiTrap Heparin HP column (Cytiva) equilibrated with the same buffer. The column matrix was washed with 10 CV heparin buffer A, followed by elution with a 0-100% linear gradient of heparin buffer B (20 mM Tris-HCl pH 8.0, 1 mM EDTA, 1 M NaCl). STAT1 and STAT1(core)^Δ183- 190, EE^ eluted at approximately 20% heparin buffer B. Fractions containing protein of interest were supplemented with TCEP (1 mM final) and concentrated on a centrifugal filter (MWCO 30,000 Da, Amicon) to 2 mL, after which the proteins were loaded on a Superdex 200 16/600 GL (Cytiva) size exclusion chromatography (SEC) column equilibrated with 20 mM Tris-HCl pH 8.0, 300 mM NaCl, 1 mM EDTA. SEC fractions corresponding to the later-eluting major peak were pooled and supplemented with TCEP (1 mM final), concentrated to ∼0.5 mM on a centrifugal filter (MWCO 30 000 Da, Amicon) and flash-frozen in liquid nitrogen.

GB1-018 and GB1-NiV-V fusions were expressed from pPEPT1 plasmids (TP, unpublished) that were transformed into *E. coli* T7 Express cells (NEB) and plated overnight on LB agar supplemented with 100 μg/mL of ampicillin. The next day transformed cells were collected and used to inoculate 1 L TB medium supplemented with 100 μg/mL of ampicillin and were grown at 37 ⁰C in 2 L flasks until OD600 of 0.8-1.2. Protein expression was induced with 0.4 mM IPTG for 3 h at 37 ⁰C. Following bacterial expression, a nickel affinity purification step was performed as described for STAT1 proteins. Fractions containing protein of interest were concentrated on a centrifugal filter (Amicon, MWCO 3000 Da) to 2 mL, after which the proteins were loaded on a Superdex 75 or Superdex 200 16/600 GL (Cytiva) SEC columns equilibrated with 20 mM Tris-HCl pH 8.0, 300 mM NaCl, 1 mM EDTA. SEC fractions corresponding to GB1 fusions were pooled, concentrated on a centrifugal filter (MWCO 3000 Da, Amicon) and flash-frozen in liquid nitrogen. For the purification of NiV-V proteins, buffers were supplemented with TCEP (1 mM final) to maintain cysteines in a reduced state.

#### Isothermal titration calorimetry (ITC)

All proteins were buffer-exchanged into ITC buffer (50 mM Tris-HCl pH 8.0, 150 mM NaCl, 1 mM EDTA, 0.1% Tween-20) using a NAP-5 size-exclusion column (Cytiva) and concentrations were determined by UV/Vis spectrophotometry and adjusted as needed. For measurements with synthetic peptides, peptides were re-suspended from lyophilised powder in MilliQ water and then concentrations were measured by UV-Vis and were adjusted to 10x the final value. Thereafter, the peptides were diluted ten-fold in ITC buffer. ITC measurements were performed on a Microcal ITC200 instrument (GE Healthcare) with 18 x 2 μL injections, 160 s interval and 5 μCal s^-1^ reference power. Baseline correction was performed using injection heats from protein-into-buffer runs. Integration of thermogram peaks and fitting of data was done using the Malvern ITC package in Origin 7.0 (Originlab). Isotherm fitting was performed using a one site model. All of the reaction conditions and fitted parameters are shown in **Table S5**.

#### Fluorescence polarisation (FP) anisotropy measurements

All proteins were buffer-exchanged into FP buffer (50 mM Tris pH 8.0, 300 mM NaCl, 1mM EDTA, 0.1% Tween-20, 1 mM TCEP) using a NAP-5 size-exclusion column (Cytiva) and concentrations were determined by UV/Vis spectrophotometry and adjusted as needed, after which, BSA was added to 0.1%. Fluorescein-conjugated pIFNGR1 12-mer peptide probe (Fluor-pIFNGR1) was first re-suspended in DMSO to 10 mM and then diluted in FP buffer plus 0.1% BSA to the required concentration. Reactions (40 μ) were set up in a 384-well non-transparent microplate (Corning, #3542). Competition reactions were performed with 10 nM Fluor-pIFNGR1 and fixed STAT1 concentration of 1.5 μM and two-fold serial dilutions of 018 or NiV-V GB1 fusions. Each dilution was measured in triplicate. Graphs show means ± SD (n=3) per dilution.

Measurements were performed on a Pherastar FS plate reader (BMG) using a FP 485/520/520 optical module. Reactions containing only 10 nM Fluor-pIFNGR1 were prepared as reference standards and were used to calibrate gain and focal height. Dose-response curves were fitted in Prism 9.0.0 (GraphPad) using a four-parameter logistic regression.

#### Peptides for ITC and FP

A 5-mer sequence (pYDKPH) of the pIFNGR1 is responsible for the vast majority of the receptor STAT1 SH2 domain interaction. For the FP assay we utilised a 12-mer peptide (5Flu- GTSFGpYDKPHVLV-NH2, PeptideSynthetics, UK) where TSFGpYDKPHVLV corresponds to 12 aa from pIFNGR1 and 5Flu-G represents an N-terminal 5-carboxyfluorescein and a spacer glycine. For our ITC measurement we utilised the 5-mer peptide (Ac-pYDKPH-NH2, Genosphere Biotechnologies) due to greater solubility compared to the 12-mer peptide. Peptides were prepared using Fmoc-based solid-phase synthesis and purity was >95% as determined by HPLC.

#### SEC-MALS

SEC-MALS was performed using a Superdex 200 Increase 10/300 column (Cytiva) equilibrated with 50 mM Tris pH 8.0, 300 mM NaCl, 1 mM EDTA, 1 mM TCEP. The column was connected to a DAWN HELEOS II light scattering detector (Wyatt Technology) and the Optilab T-rEX refractive index detector (Wyatt Technology). Scattering was detected at 664 nm wavelength at RT. One hundred μL of sample was applied at a concentration of 20 μM for STAT1 and 100 μM of GB1-018. The experimental data were recorded and processed using the ASTRA software (Wyatt Technology).

#### X-ray crystallography

A STAT1 core fragment crystallography construct (STAT1^136-684,Δ183-190,H182A,E393A,E394A^) was prepared harbouring a loop deletion at the apex of the coiled coil domain (Δ183-190,H182A) and surface entropy-reducing mutations (E393A,E394A). The STAT1-018 complex was co- crystallised using sitting-drop vapour diffusion in a 96-well MRC plate format. The complex was prepared by mixing STAT1^136-684,Δ183-190,H182A,E393A,E394A^ in SEC buffer (20 mM Tris-HCl pH 8.0, 300 mM NaCl, 1 mM EDTA) and 018 21-mer peptide (Ac- MWSVFIHGHDGSNKGSKTYTS-NH2, Genosphere Biotechnologies) in 20 mM Tris-HCl pH 8.0, to a final concentration of 5 mg/mL protein and 2 mg/mL peptide. Three hundred nL of the complex was mixed with 300 nL of the crystallisation condition using a Mosquito liquid handling robot (TTP Labtech). Crystals were obtained using the following condition: 16% PEG 3350, 175 mM KCl, 125 mM (NH4)2SO4. Cryoprotectant solution containing the crystallisation condition and 30% ethylene glycol was added to the drop and crystals were incubated for 1 min. A crystal was then harvested and cryo-cooled in liquid nitrogen. Diffraction data were collected at Diamond Light Source (Harwell, UK) synchrotron radiation source, beamline i04. Diffraction images were processed with autoPROC (Vonrhein et al., 2011). Molecular replacement phasing was used with STAT1 core residues 133-683 (PDB ID: 1YVL) as a search model. The structure was refined without peptide first and the peptide was built into the clearly visible electron density manually **(Figure S6A)**. Manual real-space refinement was done in Coot (Emsley et al., 2010) and automated refinement with phenix.refine (Liebschner et al., 2019) and autoBUSTER (Smart et al., 2012). Crystallographic data and refinement statistics are shown in **Table S4**. The coordinates and corresponding structure factors have been deposited to the PDB under accession number PDB: 7nuf.

## Supplemental information

**Table S1.**
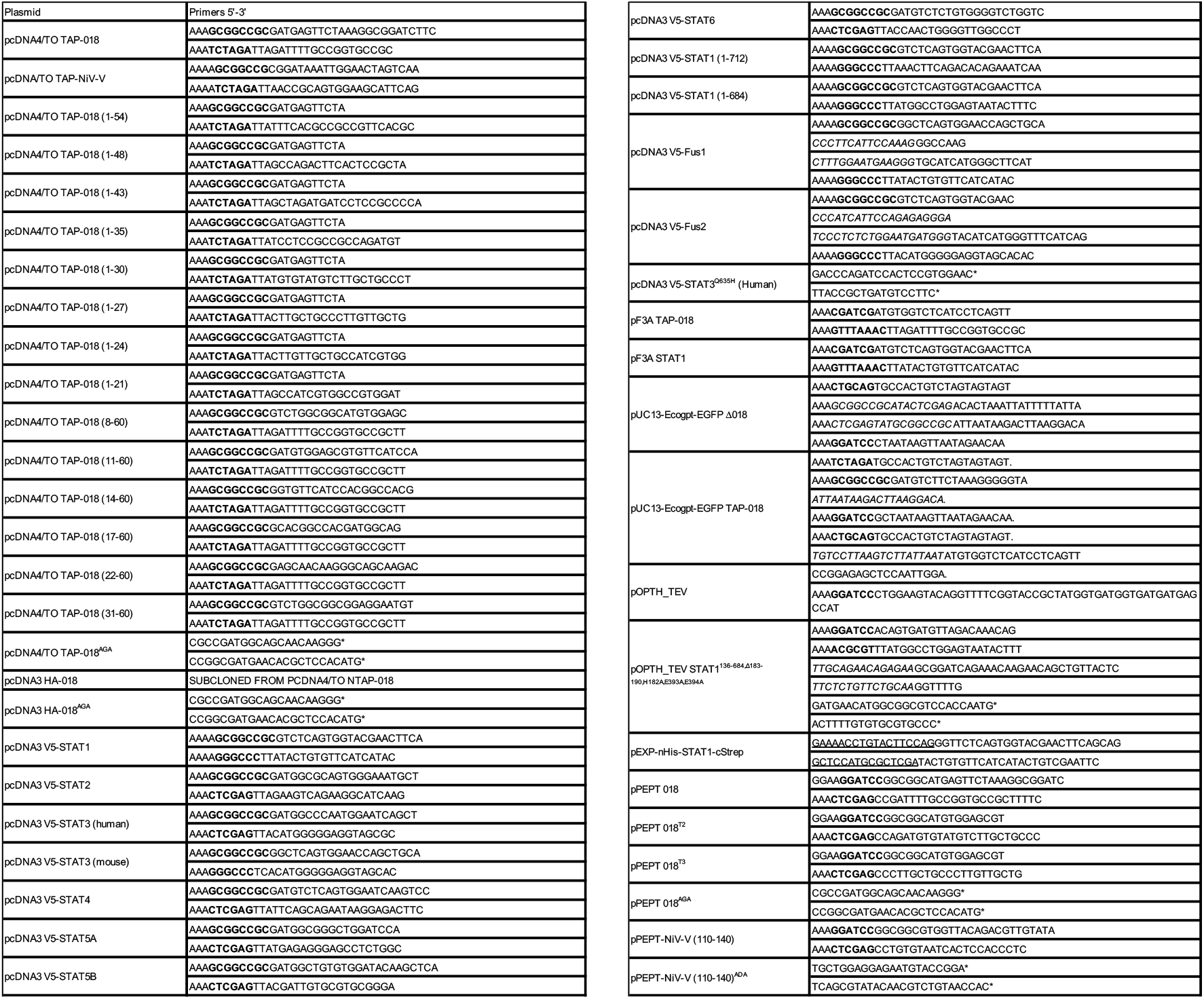
Oligonucleotide primers for construction of recombinant DNA. Related to STAR methods. Cloning sites are highlighted in bold (for restriction digest cloning) or underlined (for ligation- independent cloning). Complementary sequences for overlapping PCR are italicised. Site- directed mutagenesis primers are indicated with an asterisk (*). All V5 and TAP-tagged proteins are tagged at the N terminus.

**Table S2.**
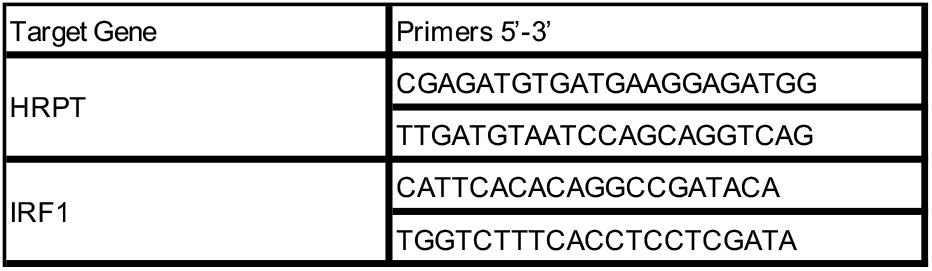
Oligonucleotide primers for RT-qPCR. Related to STAR methods.

**Table S3.**
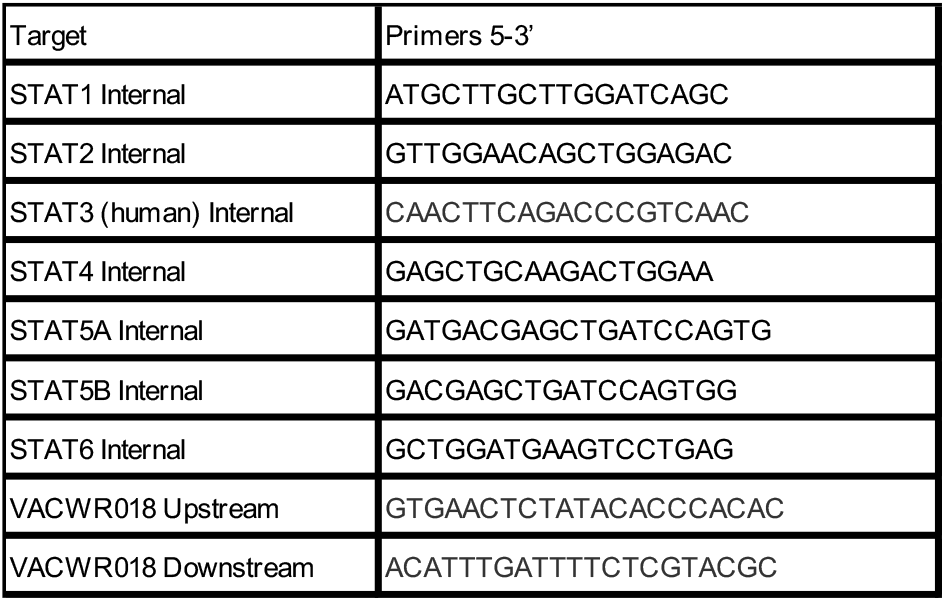
Oligonucleotide primers for analytical PCR or sequencing. Related to STAR methods.

**Table S4.**
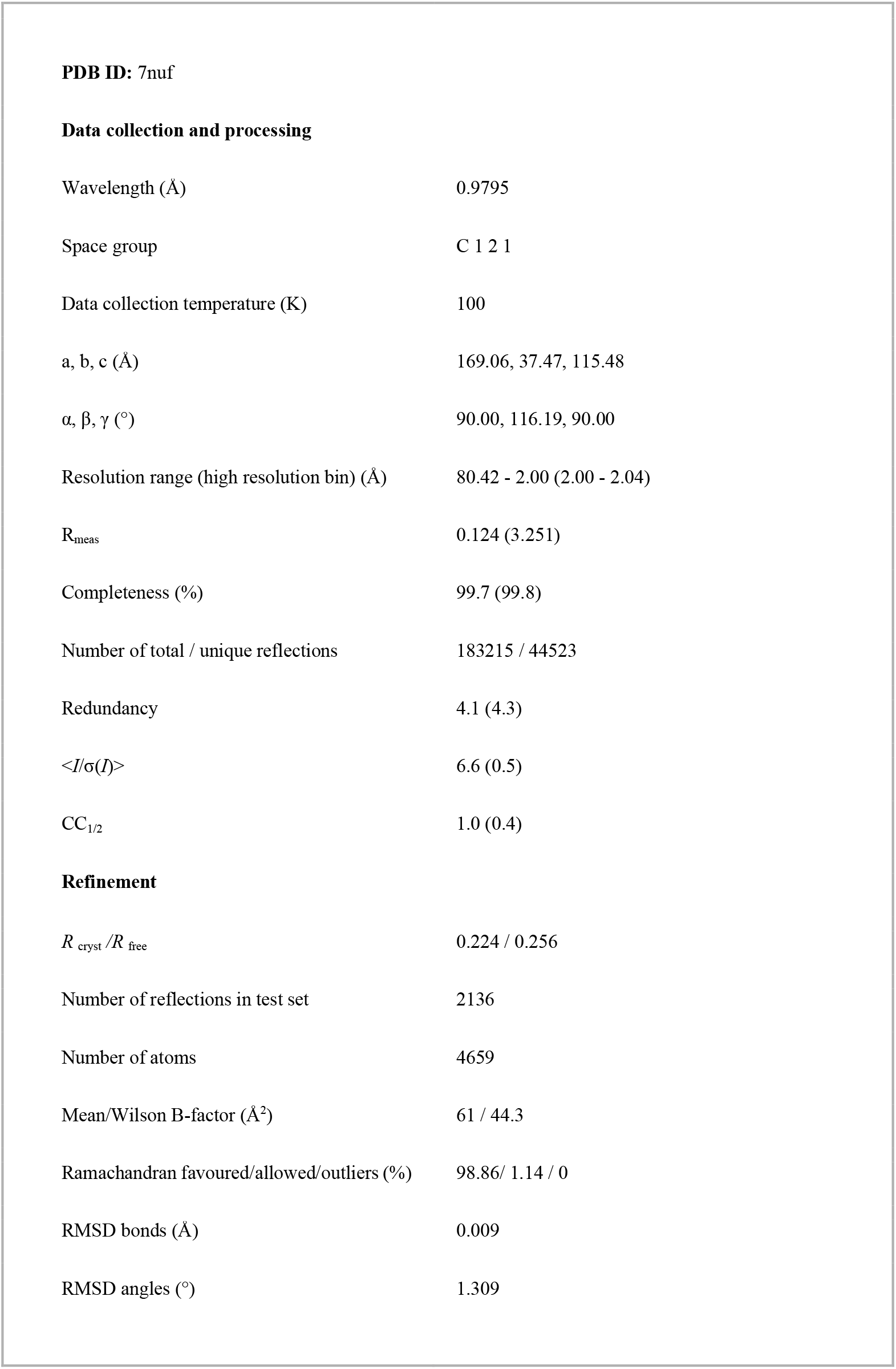
X-ray crystallographic data collection and refinement statistics.

**Data collection and processing**
|  |  |
| --- | --- |
| Wavelength (Å) | 0.9795 |
| Space group | C 1 2 1 |
| Data collection temperature (K) | 100 |
| a, b, c (Å) | 169.06, 37.47, 115.48 |
| $\alpha$, $\beta$, $\gamma$ (°) | 90.00, 116.19, 90.00 |
| Resolution range (high resolution bin) (Å) | 80.42 - 2.00 (2.00 - 2.04) |
| $R_{\text{meas}}$ | 0.124 (3.251) |
| Completeness (%) | 99.7 (99.8) |
| Number of total / unique reflections | 183215 / 44523 |
| Redundancy | 4.1 (4.3) |
| $\langle I/\sigma(I) \rangle$ | 6.6 (0.5) |
| $CC_{1/2}$ | 1.0 (0.4) |
| **Refinement** |  |
| $R_{\text{cryst}}/R_{\text{free}}$ | 0.224 / 0.256 |
| Number of reflections in test set | 2136 |
| Number of atoms | 4659 |
| Mean/Wilson B-factor (Å<sup>2</sup>) | 61 / 44.3 |
| Ramachandran favoured/allowed/outliers (%) | 98.86/ 1.14 / 0 |
| RMSD bonds (Å) | 0.009 |
| RMSD angles (°) | 1.309 |

**Table S5.** ITC experimental conditions and fitted parameters.

| Reaction | Cell | Syringe | $K_d$, nM | N | $\Delta H$, cal / mol | $\Delta S$, cal / mol / deg |
| --- | --- | --- | --- | --- | --- | --- |
| 018 + STAT1 | 10 μM STAT1 | 100 μM 018 | 291 | 1.02 | $-1.93 \times 10^4$ | -34.9 |
| 018T2 + STAT1 | 15 μM STAT1 | 150 μM 018T2 | 235 | 1.01 | $-1.87 \times 10^4$ | -32.3 |
| 018T3 + STAT1 | 15 μM STAT1 | 350 μM 018T3 | $>10^4$ | 1.00 (fixed) | $-2.01 \times 10^4$ | -46.8 |
| 018AGA + STAT1 | 10 μM STAT1 | 100 μM 018AGA | No binding | n.a. | n.a. | n.a. |
| pIFNGR1 + STAT1 | 10 μM STAT1 | 300 μM pIFNGR1 5-mer | $7.6 \times 10^3$ | 0.708 | -9172 | -7.33 |
| pIFNGR1 + STAT1 / 018 | 10 μM STAT1 + 50 μM 018 | 300 μM pIFNGR1 5-mer | No binding | n.a. | n.a. | n.a. |
| pIFNGR1 + STAT1 / 018 AGA | 10 μM STAT1 + 50 μM 018 AGA | 300 μM pIFNGR1 5-mer | $7.8 \times 10^3$ | 0.79 | -8001 | -3.48 |
| pIFNGR1 + STAT1 / Niv-V | 10 μM STAT1 + 200 μM Ni V | 300 μM pIFNGR1 5-mer | No binding | n.a. | n.a. | n.a. |
| pIFNGR1 + STAT1 / Niv-V ADA | 10 μM STAT1 + 200 μM Ni V ADA | 300 μM pIFNGR1 5-mer | $7.2 \times 10^3$ | 0.88 | -8125 | -3.72 |

**Figure S1.**
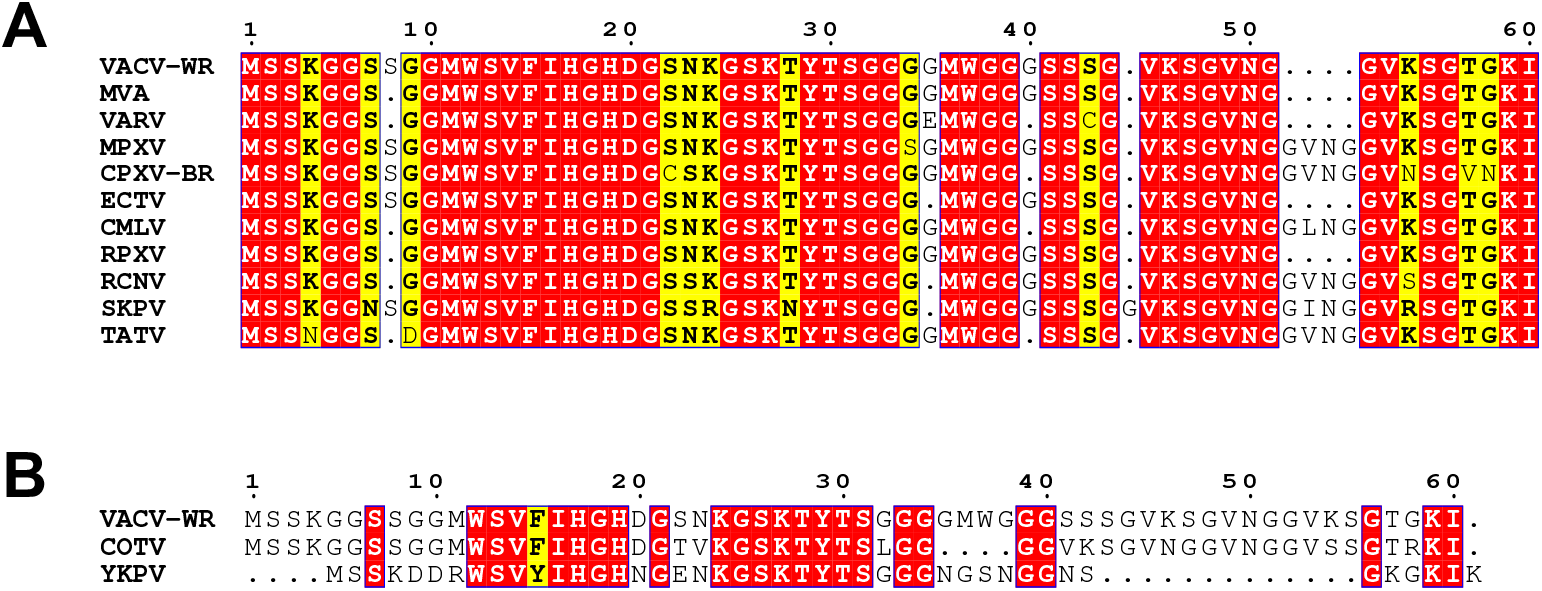
Sequence alignment of 018 orthologues. (**A**) Alignment of 018 orthologues from representative orthopoxviruses: vaccinia strain Western Reserve (VACV-WR), modified vaccinia Ankara (MVA), variola virus (VARV), monkeypox virus (MPXV), cowpox virus strain Brighton Red (CPXV), ectromelia virus (ECTV), camelpox virus (CMPV), rabbitpox virus (RPXV), racconpox virus (RCNV), skunxpox virus (SKPV) and taterpox virus (TATV). The 018 ORF is absent from VACV strain Copenhagen. (**B**) Alignment of 018 poxvirus orthologues from Cotia virus (COTV) and yokapoxvirus (YKPV), which sit outside of the orthopoxvirus genus. Identical residues are shown in red, similar residues are shown in yellow (**A-B**).

**Figure S2.**
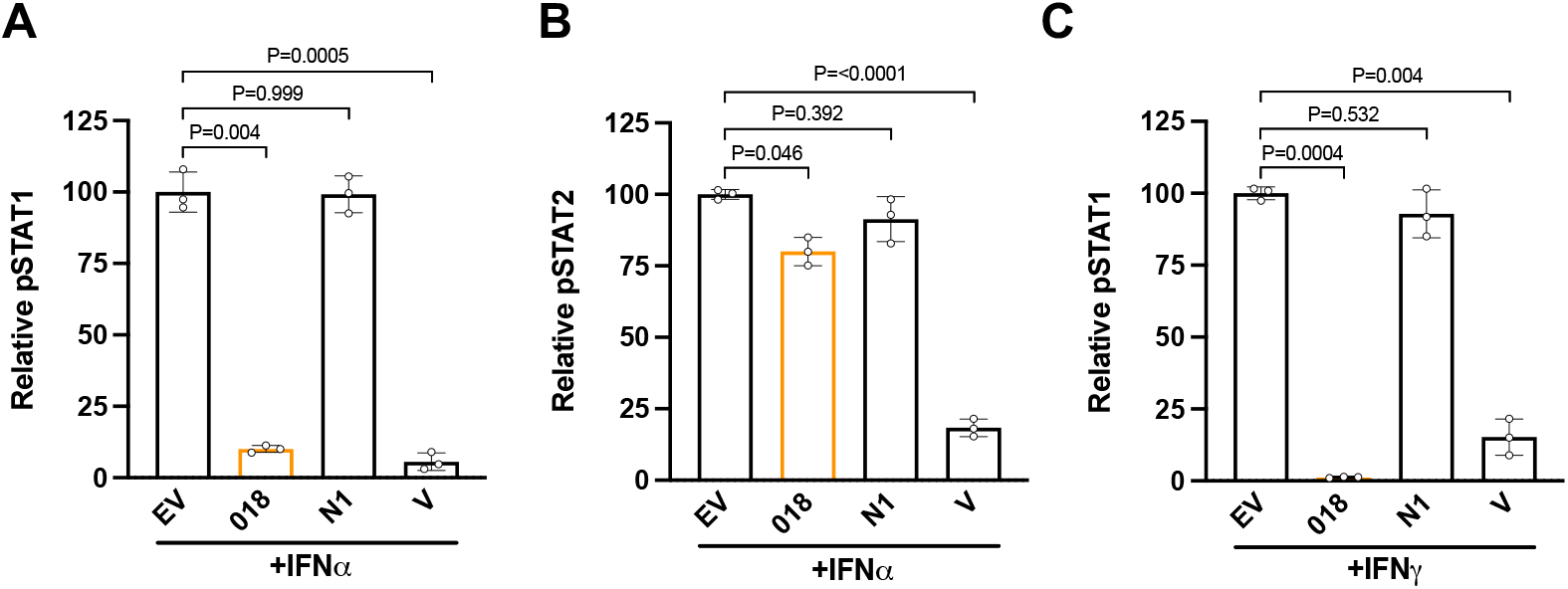
Related to Figure 2. Quantification of relative band intensities (**Figure 2**) for pSTAT1 **(A)** and pSTAT2 (**B**) from (**Figure 2C)** and pSTAT1 **(C)** from (**Figure 2D**). pSTAT levels were normalised against total STAT levels and made relative to EV condition. Means ± SD (n=3 per condition) are shown. Significances were calculated by Dunnett’s T3 multiple comparisons test.

**Figure S3.**
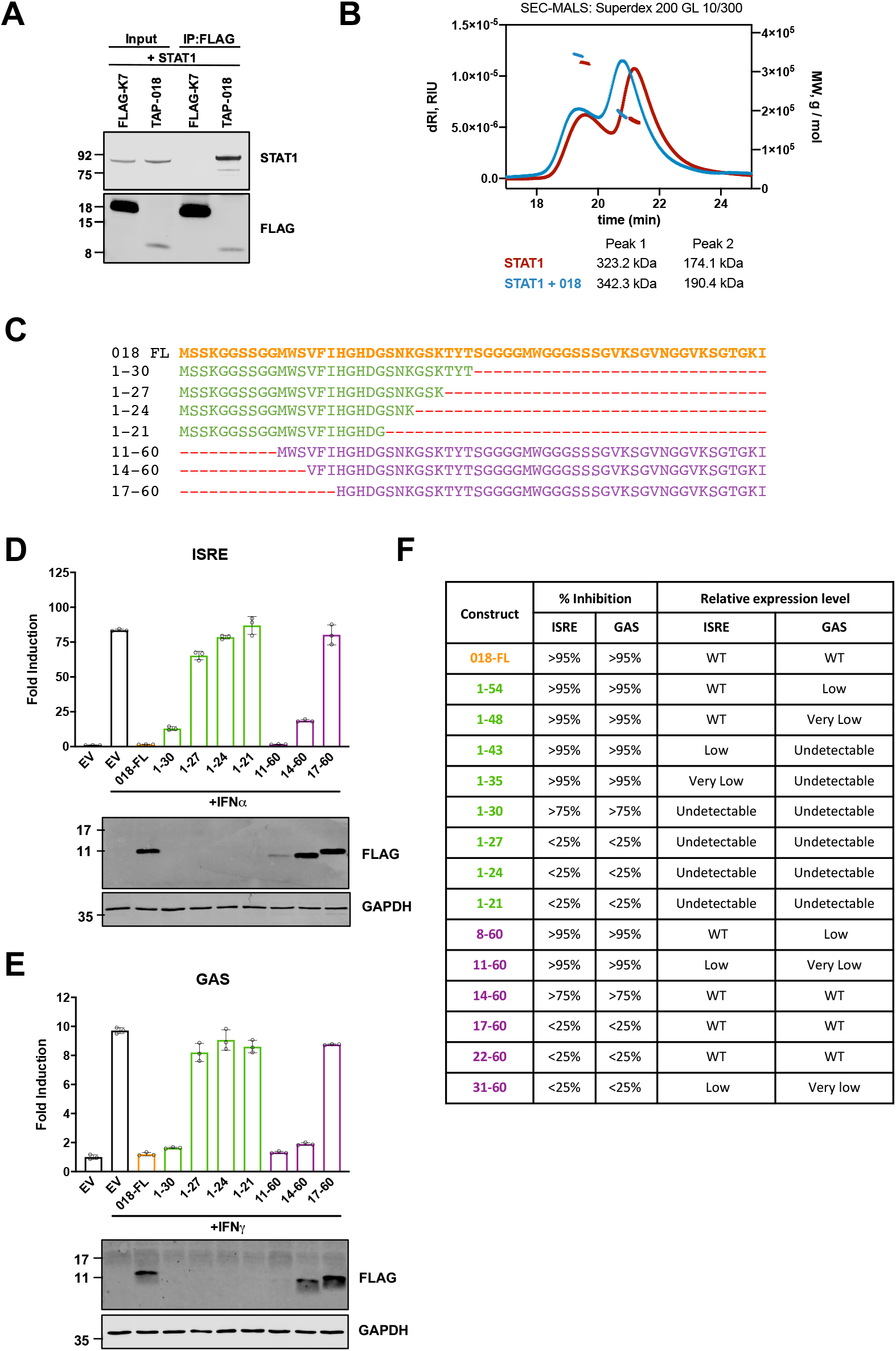
Related to Figure 3. (**A**) 018:STAT1 interaction using a cell-free transcriptional and translation system. Untagged- STAT1 was co-expressed along with either FLAG-tagged K7 or TAP-tagged 018 using a wheat germ cell free transcriptional and translation system. FLAG-K7 and TAP-018 were precipitated using M2-FLAG affinity gel and purified proteins were analysed by immunoblotting with α- FLAG and α-STAT1 antibodies. (**B**) SEC-MALS measurements of purified free STAT1 (red) and STAT1:GB1-018 complex (blue). One hundred μl samples were loaded on a Superdex 200 GL 10/300 column and scattering and refractive index of the eluting peaks were measured. Concentration of 20 μM for STAT1 and 100 μM of GB1-018 were applied. (**C**) Sequences for C-terminal (green) and N-terminal (purple) 018 refined truncation mutants. (**D**) HEK 293T cells or (**E**) HeLa cells were transfected with reporter plasmids ISRE-Luc (**D**) or GAS-Luc (**E**) plus *TK*-*Renilla* and vectors expressing 018 truncation mutants from (**C**) fused to a TAP-tag. Cells were stimulated with IFNα (1000 U/mL) (**D**), or IFNγ (25 ng/mL) (**E**) for 6 h (**D**) or 8 h (**E**) and luciferase values were measured. Means ± SD (n=3 per condition) are shown. Lysates were prepared and analysed by immunoblotting with α-FLAG and α-GAPDH. (**F**) Summary table of all C-terminal (green) and N-terminal (purple) 018 truncation mutants describing the percentage inhibitory activity (>95%, >75% (but less then >95%) or <25%) and relative protein expression levels (wild type (WT), low, very low or undetectable) for ISRE (IFNα) and GAS (IFNγ) reporters. Data from (**D-E**) are representative of 2 individual experiments.

**Figure S4.**
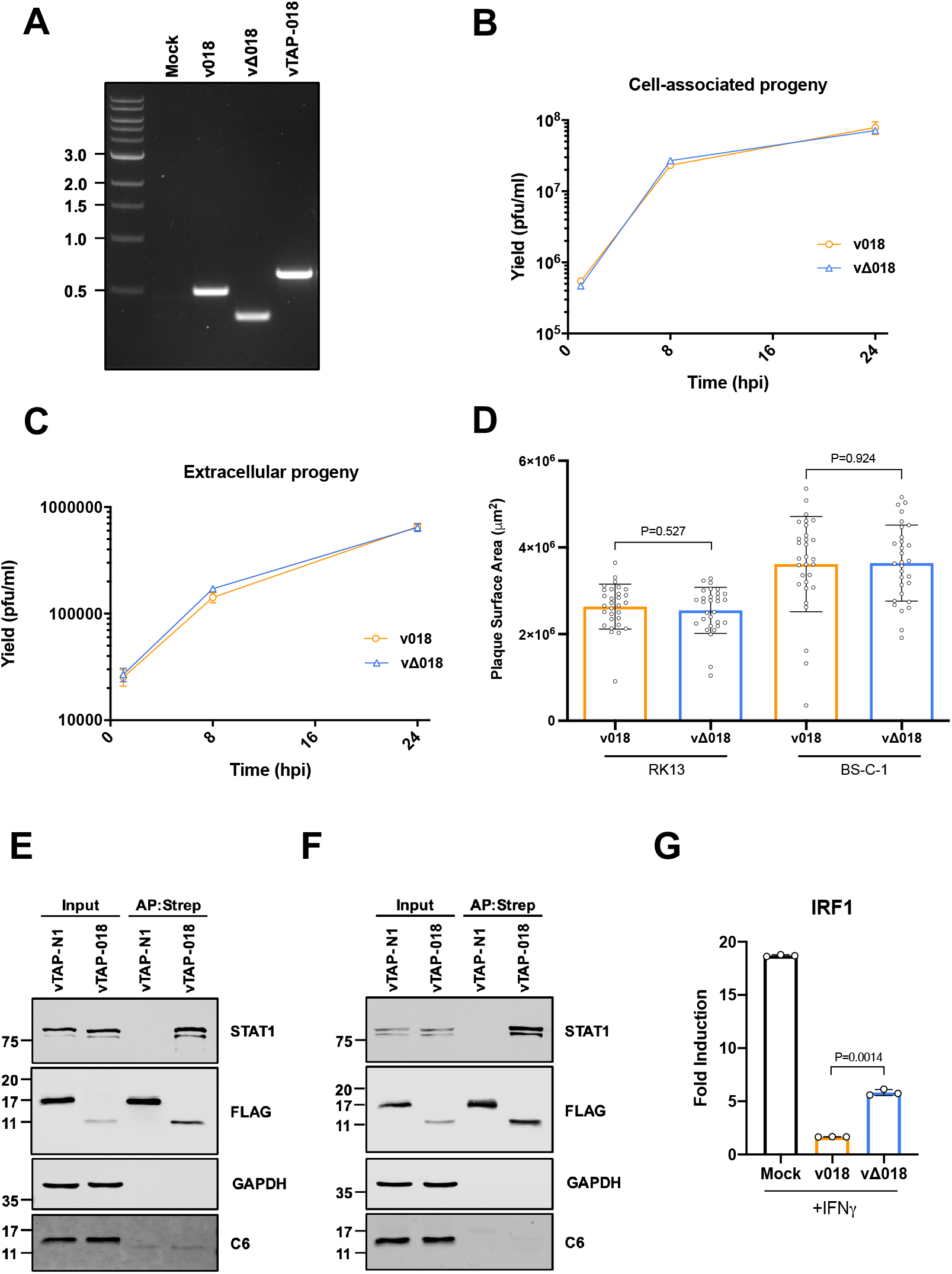
Related to Figure 4. (**A**) PCR amplification of genomic DNA from the indicated viruses using primers upstream and downstream of the 018 ORF. The position of DNA size markers (kb) are shown on the left side of the image. (**B, C**) BS-C-1 cells were infected with either v018 (orange) or vΔ018 (blue) at 5 pfu/cell. At 1, 8 and 24 h p.i. infectious virus titres associated with cells (**B**) and in the supernatants (**C**) was determined by plaque assay on BS-C-1 cells. Means ± SD (n=2 per condition) are shown and P=>0.05 for all timepoints. **(D)** RK13 or BS-C-1 cells were infected at 30 pfu per well with either v018 (orange) or vΔ018 (blue). At 72 h p.i. monolayers were stained and plaque surface areas were quantified. Means ± SD (n=30 plaques per condition) are shown. **(E)** BS-C-1 cells or **(F)** MEFs were infected with vTAP-018 or vTAP-N1 at 5 pfu/cell for 12 h. TAP-tagged proteins were affinity-purified by Strep-Tactin and whole cell lysates (Input) and affinity-purified proteins (AP:Strep) were analysed by immunoblotting with α-FLAG, α-GAPDH, α-STAT1 and α-C6. Data for (**B-F**) are representative of two individual experiments. (**G**) A549 cells were mock-infected or infected with v018 or vΔ018 at 10 pfu/cell. At 2 h p.i. cells were stimulated IFNγ (25 ng/mL) for 1 h. Total RNA was extracted and mRNA for IRF1 was analysed by RT-qPCR. Means ± SD (n=3 per condition) are shown. Data are representative of two individual repeats. Significances were determined using Unpaired t-test with Welch’s correction (**B-D**, **G**).

**Figure S5.**
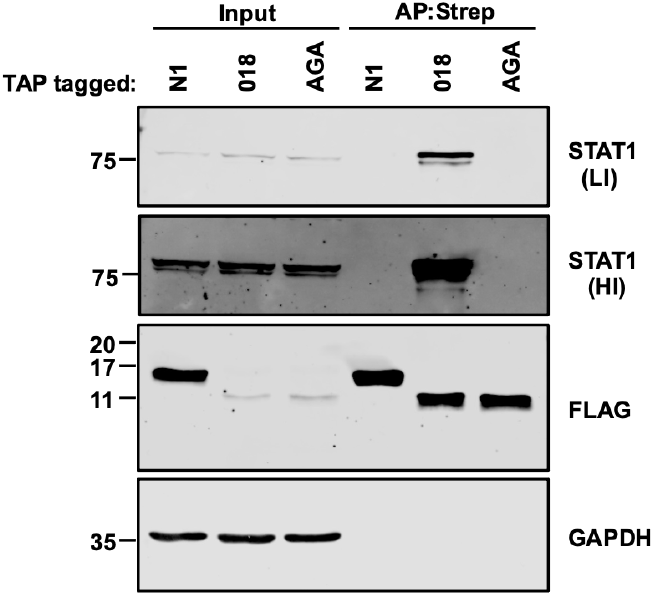
Related to Figure 6. TAP-tagged 018, 018 AGA (labeled AGA) and N1 were expressed by transfection in 2fTGH cells and affinity purified by Strep-Tactin. Whole cell lysates (Input) and affinity-purified proteins (AP:Strep) were analysed by immunoblotting with α-FLAG, α-GAPDH and α- STAT1. A high intensity (HI) and low intensity (LI) scan of α-STAT1 are shown.

**Figure S6.**
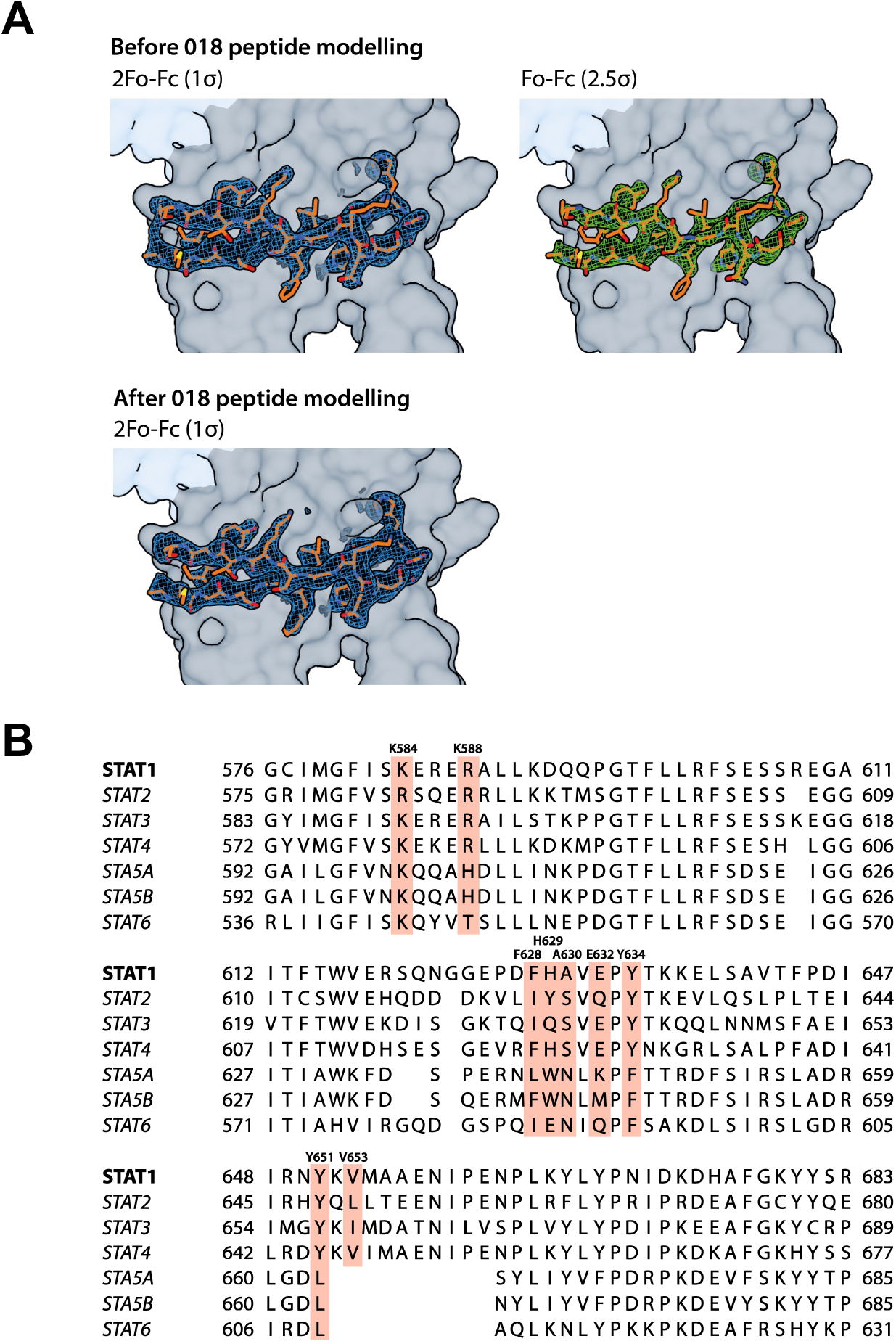
Related to Figure 7. **(A)** Electron density maps of 018 peptide before and after peptide modelling and refinement. **(B)** Alignment of STAT family SH2 domains with residues that form contacts with 018 in the 018:STAT1 crystal structure highlighted in orange.

**Figure S7.**
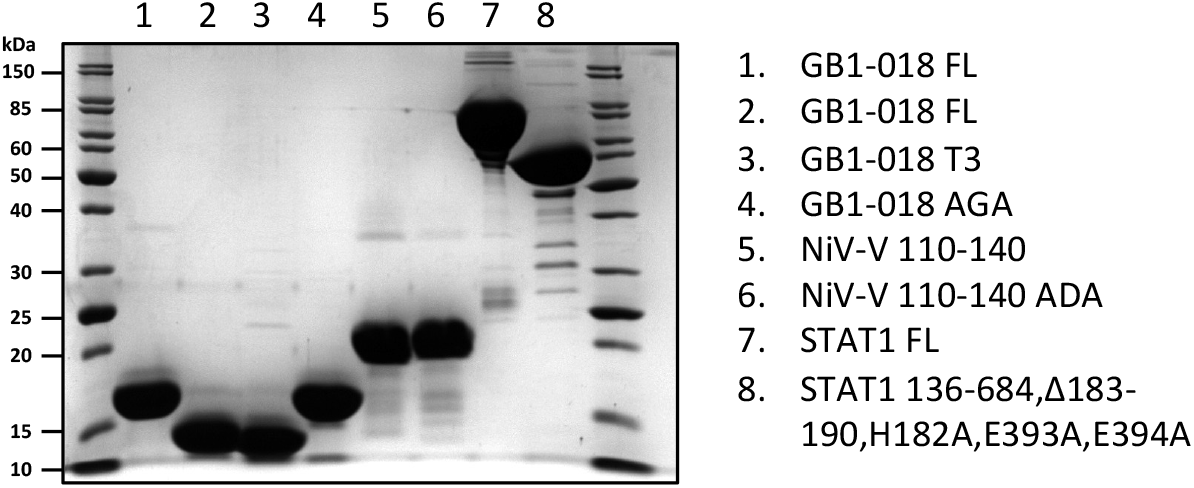
Coomassie-stained SDS-polyacrylamide gel of purified recombinant proteins.

